# Sequence and Structural Alignments Reveal Insights into ANKLE2 Evolution and Function

**DOI:** 10.64898/2026.01.21.700886

**Authors:** Adam T Fishburn, Cole J Florio, Chase L S Skawinski, Sydney S Becker, Ethan Holleman, Avery E Robertson, Rees Sitchon, Frédéric Chédin, Priya S Shah

## Abstract

ANKLE2 is an enigmatic protein with emerging roles in cell division, development, and virus replication. While ANKLE2 orthologs are present in all animals, its domain composition has evolved over time. ANKLE2’s two namesake domains, the ankyrin repeat and LEM domains, have clear and defined roles; however nearly all ANKLE2 orthologs have at least three other structured domains with poorly understood purposes. In this study, we performed sequence and structural alignments of ANKLE2 orthologs to improve our understanding of the protein’s evolution and function. We identified that ANKLE2’s transmembrane domain likely evolved more recently and coincided with loss of VAPA interaction as a membrane anchoring mechanism. We show that despite stark differences in amino acid sequence, the structure of the LEM and ankyrin repeat domains are highly conserved across ANKLE2 orthologs. To investigate ANKLE2’s uncharacterized domains, we performed structural alignments to identify similar proteins. This revealed surprising similarities between portions of ANKLE2 and nuclease or nucleic acid-binding proteins. However, ANKLE2 lacks key motifs imparting function in these domains, which was confirmed by experimental interrogation. We further identified that loss of ANKLE2 is correlated with changes in DNA damage response and micronuclei formation. We believe this methodology demonstrates the power of combining structural predictions with classical molecular techniques in exploring poorly understood proteins.

**Importance:** ANKLE2 is a scaffolding protein present in all animals; however much of its function is poorly understood. By evaluating ANKLE2 sequence and structure from many different organisms and comparing its various domains with other proteins, we gain insight into how ANKLE2 evolved and what cellular roles it might be fulfilling. Further, this approach can be used to investigate other understudied or uncharacterized proteins.

## 1 INTRODUCTION

One fundamental philosophy of molecular biology is that structure dictates function. However, we often determine how similar proteins are by their relation at the sequence level. Recent advancements in protein structural prediction now allow us to better compare orthologous proteins across the tree of life. Further, we can apply these techniques to shed light on the purpose and function of uncharacterized domains within understudied proteins. We sought to apply this approach to ankyrin repeat and LEM domain containing protein 2 (ANKLE2), a scaffolding protein with known roles in cell division and brain development, but with broader functions still unknown (Fishburn et al. 2024).

ANKLE2 was first described in 2004 as one of the few LEM domain proteins (Lee and Wilson 2004), which mediate interaction with barrier to autointegration factor (BAF) at the inner nuclear lamina (Shumaker et al. 2001). Here, ANKLE2 coordinates the phosphorylation state of BAF by facilitating interaction with the kinase VRK1 and PP2A phosphatase complex (Asencio et al. 2012). Phosphorylation of BAF is a key trigger in the disassembly of the nuclear envelope in early mitosis, and dephosphorylation is required for proper nuclear reassembly after division (Asencio et al. 2012; Snyers et al. 2018). ANKLE2 is also critical for proper brain development. In humans, certain mutations in *ANKLE2* are associated with congenital primary microcephaly, a condition in which the brain is not fully developed at birth (Yamamoto et al. 2014; Link et al. 2019; Thomas et al. 2022). In fruit fly models, depletion or mutation of *Ankle2* results in abnormal nuclear envelope morphology and reduced brain size of third instar larva (Yamamoto et al. 2014; Shah et al. 2018; Link et al. 2019). This microcephaly-like phenotype arises from underlying defects in the asymmetric division of neuroprogenitor cells (Link et al. 2019). In zebrafish, loss of *ankle2* causes progressive microcephaly and defects in spermatogenesis (Apridita Sebastian et al. 2022). Importantly, fly phenotypes can be rescued by overexpression of the human ANKLE2 ortholog, supporting that ANKLE2’s function in mitosis and brain development is broadly conserved across evolutionarily distant organisms (Yamamoto et al. 2014). While ANKLE2 certainly has critical roles in certain points of mitosis and neurodevelopment, it maintains consistent expression throughout the cell cycle, the body, and development (Fagerberg et al. 2014; Cardoso-Moreira et al. 2019), raising questions about its broader functions in cell biology and physiology. During interphase, ANKLE2 is localized to the endoplasmic reticulum (ER), however, its function in this organelle is largely unexplored. ANKLE2 is known to interact with a variety of other host proteins from large-scale proteomic screens (Hein et al. 2015; Huttlin et al. 2021), and it is speculated that its interaction with PP2A may regulate the phosphorylation of other substrates beyond BAF. Our previous work characterized the organization of ANKLE2 orthologs from several model organisms where we observed that key domains are not present in all species (Fishburn et al. 2024). These observations draw into question the true nature of ANKLE2 function, the role these domains play in those functions, and how these domains emerged across evolutionary time.

In this study, we explore ANKLE2 function using an evolutionary and structural perspective. We examine ANKLE2 orthologs across the animal kingdom to first gain insight into each domain’s level of conservation and then draw conclusions about the protein’s evolutionary history. To validate some of our hypotheses, we characterized a number of ANKLE2 orthologs experimentally. While these orthologs share expectedly low amino acid identity as phylogenetic distance grows, we found that the structural similarity of each domain is highly conserved, even between evolutionarily distant orthologs. To probe the identity and potential function of several uncharacterized ANKLE2 domains, we employed this structure-based approach to identify proteins that contain the most similar sub-structures. This revealed striking similarities between portions of ANKLE2 and GIY-YIG endonucleases or R-loop binding ribonucleases.

However, despite these structural similarities, our experiments revealed that ANKLE2 did not share these nuclease functions, likely because of differences in key motifs that impart nuclease activity and nucleic acid binding. Finally, we identify a previously unknown role for ANKLE2 in regulating the response to DNA damage. Together, our work demonstrates novel aspects of ANKLE2 function derived from an evolutionary and structural approach.

## 2 RESULTS

### 2.1 Evaluation of ANKLE2 domain organization across metazoa

ANKLE2 is named after the LEM and ankyrin repeat domains it contains. However, human ANKLE2 structural predictions contain four additional distinct regions with high confidence (Figure 1). The N-terminal transmembrane (TM) domain is consistently predicted by TM prediction algorithms such as DeepTMHMM (Figure 1) (Hallgren et al. 2022) and mediates localization to the ER (Elkhatib et al. 2017; Saeng-chuto et al. 2025). After the TM is the LEM domain, a ∼50 amino acid three-helical domain, which mediates interaction with BAF. Following the LEM domain is a peculiar structure occasionally annotated as “Caulimovirus viroplasmin” or “Caulimovirus domain (CD)” which is named after viroplasmin proteins found in Revtraviricetes retroviruses (Mushegian et al. 1994; Cerritelli et al. 1998). Mechanistic study of this region in ANKLE2 has been extremely limited but is important for interaction with PP2A subunits (Asencio et al. 2012; Saeng-chuto et al. 2025). After the CD is the ankyrin repeat domain (ANK) which is thought to contribute to the broader scaffolding function of ANKLE2, however this has never been explicitly determined. Following the ankyrin repeat domain are two distinct structures that are entirely uncharacterized (Unc-1 and Unc-2). However, mutations in these regions are associated with congenital disease, suggesting they serve some crucial role in ANKLE2 function (Fishburn et al. 2024). Despite these structured domains, the numerous intrinsically disordered regions in ANKLE2 have complicated experimental determination of its full structure.

**Figure 1:**
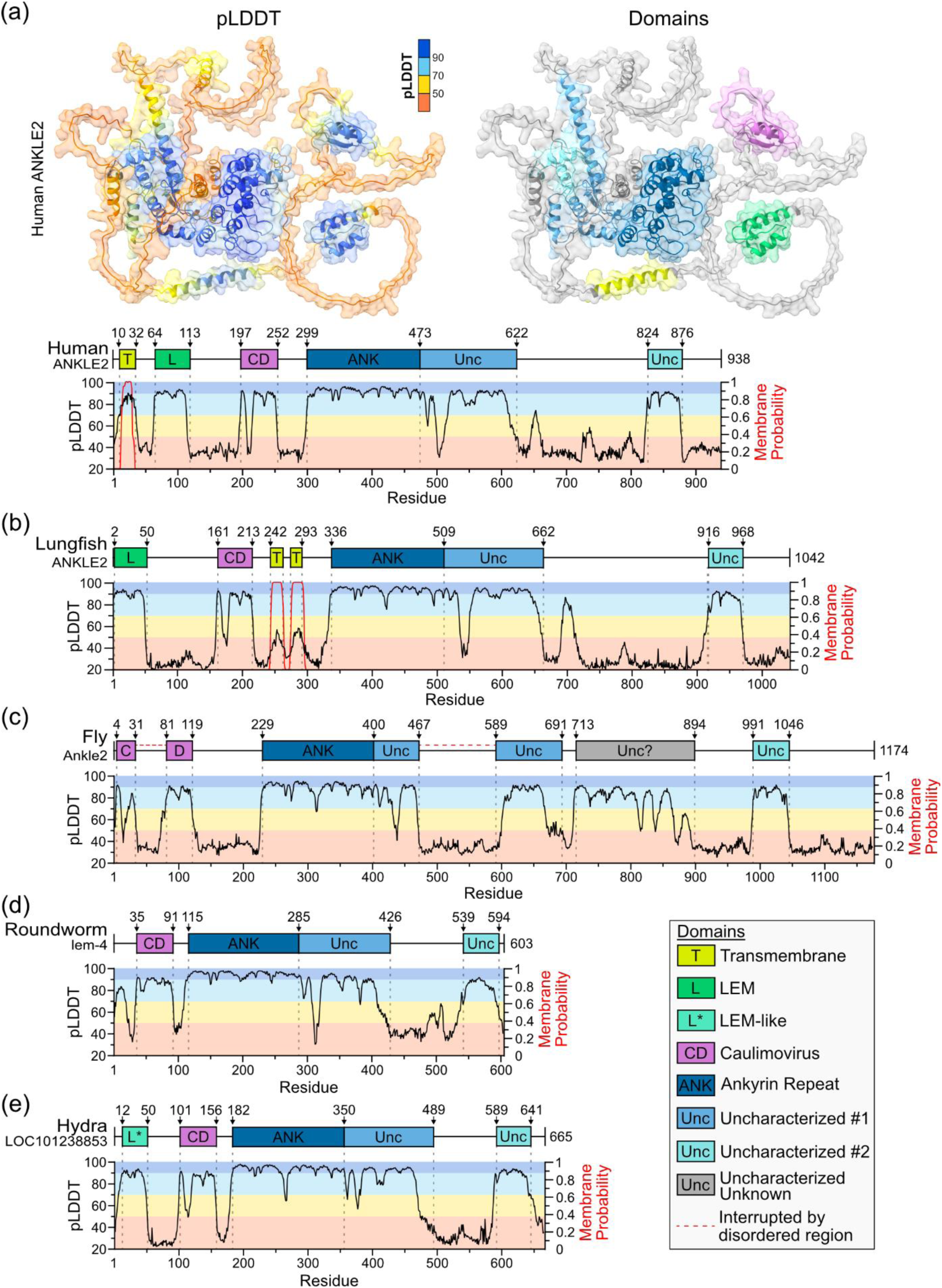
AlphaFold as a predictive tool for studying ANKLE2. (a-e) ANKLE2 ortholog protein maps showing AlphaFold pLDDT and DeepTMHMM transmembrane probability distribution.

Experimental studies on ANKLE2 have been limited to humans (ANKLE2), zebrafish (*D. rerio,* Ankle2), fruit flies (*D*. *melanogaster,* Ankle2), and the roundworm (*C. elegans,* lem-4). We previously mapped the organization of these ANKLE2 orthologs to briefly explore the conservation across these species (Fishburn et al. 2024). Here, we extended this approach to more thoroughly cover orthologs across the Metazoan kingdom. Using NCBI’s Gene search engine and the AlphaFold2 database (Mirdita et al. 2022), we identified and evaluated the predicted structure of 31 total ANKLE2 orthologs spanning 11 classes within Chordata and 10 additional metazoan phyla. Species were chosen to provide as expansive coverage as possible, though some classes/phyla could not be included due to lack of a sequenced genome.

First, to compare all these sequences against each other and identify orthologous domains, we performed a multiple sequence alignment (MSA) using ClustalOmega (Madeira et al. 2024) (Figure S2). Unsurprisingly, there was the highest amino acid sequence identity between Chordate orthologs, with >50% identity between sequences until hagfish. After this, sequence identity is consistently between ∼25-35% which is considered only low to moderate conservation. We combined our MSA results with predicted structures to annotate well-defined structured domains in ANKLE2 orthologs. We designed a non-quantitative cladogram of these orthologs and their domain composition, based on established animal phylogeny (Dunn et al. 2014; Telford et al. 2015), which will serve as a key point of reference throughout this study (Figure 2).

**Figure 2:**
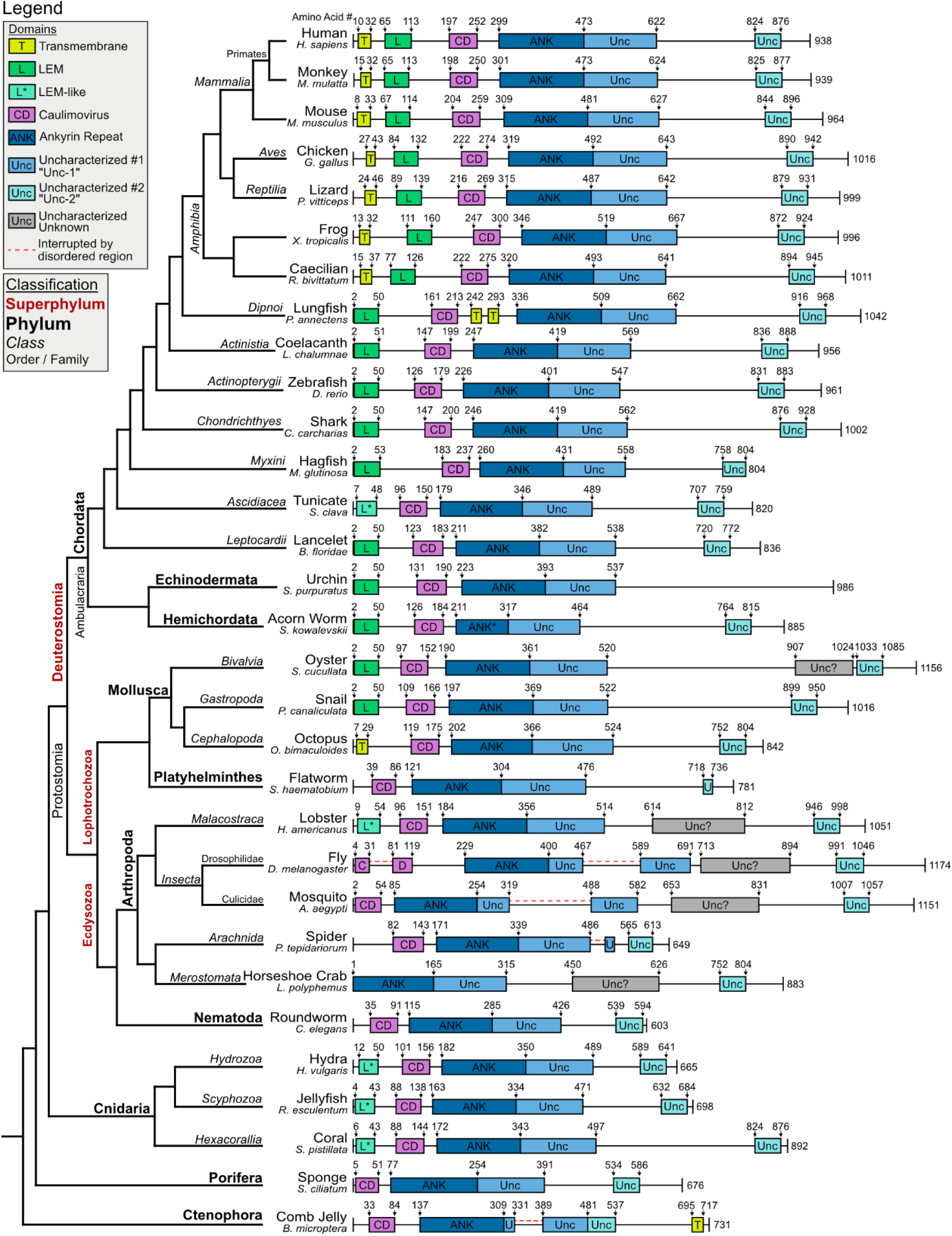
Cladogram of ANKLE2 protein composition across the animal kingdom. Presumptive ANKLE2 ortholog sequences were acquired from NCBI, and AlphaFold was used to predict the structure. A combination of AlphaFold confidence and MUSCLE sequence alignment was used in determining domain start and endpoints. Cladogram was drawn using established phylogenic relationships (Dunn et al. 2014; Telford et al. 2015).

### 2.2 Expression of ANKLE2 orthologs reveal trends in localization and interaction evolution

The first pattern we observed across ANKLE2 orthologs pertained to the presence and location of likely TM domains, which were determined using DeepTMHMM (Figures 1 and 2) (Hallgren et al. 2022). N-terminal TM domains are clearly present from Mammalia through Amphibia but are seemingly absent starting at the lobe-finned coelacanth (Figure 2). Interestingly, in-between these are lungfish, which contain two neighboring TM domains between CD and ANK, which would retain overall membrane topology with all functional domains on the cytoplasmic side of the ER-membrane (Figure 1). This observation suggests that the development of the TM domain, as a means to mediate ER-localization, is a relatively recent evolutionary event. This is supported by recent evidence that fruit fly Ankle2, which lacks a TM, mediates its ER-localization through a protein interaction with Vap33 (Li et al. 2025).

We hypothesized that all ANKLE2 orthologs with TM-domains are directly ER-associated, and those without TM-domains mediate this localization through interaction with tethering proteins homologous to Vap33. To test this hypothesis, we determined if various ANKLE2 orthologs interact with VAPA, the human ortholog of fly Vap33. We expressed human ANKLE2 and seven other animal orthologs with C-terminal 3xFLAG tags for purification and visualization (Figure 3a). We chose these specific orthologs based on the most used model organisms and to capture the variability in ANKLE2 architecture. We transfected these into HEK293T cells and performed affinity-purification to co-purify interacting proteins (Figure 3b). As predicted, human and mouse ANKLE2 did not interact with VAPA. However, five orthologs without the TM did, supporting that this interaction evolved to mediate ER-localization and was lost once TM-domains arose. Interestingly, the *Hydra* ANKLE2 ortholog did not interact with VAPA, which we attribute to the lack of a clear VAPA/Vap33 ortholog in *Hydra*. This suggests the interaction was an early evolutionary development in animal ANKLE2.

**Figure 3:**
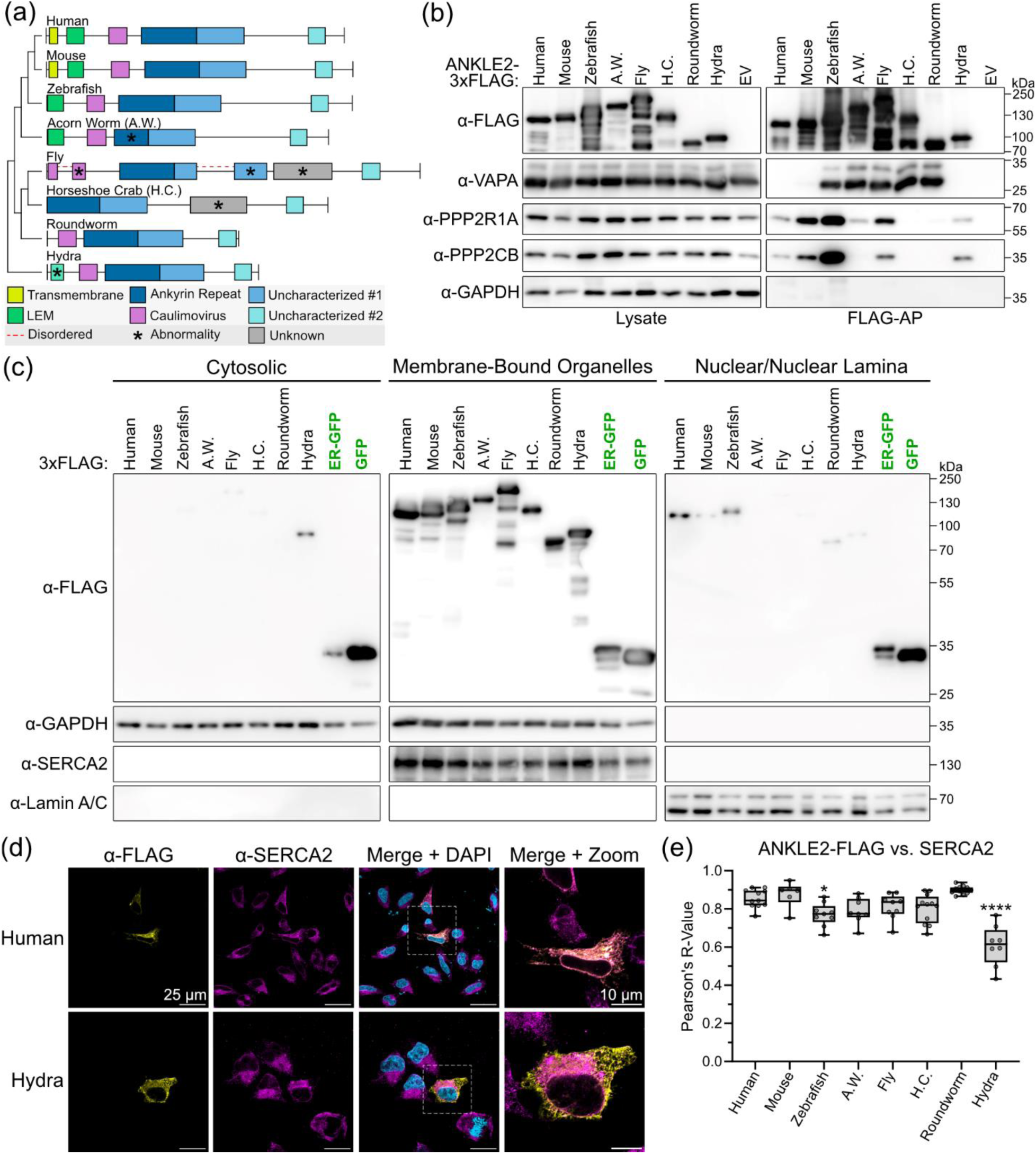
Characterization of ANKLE2 ortholog protein-interactions and subcellular localization. (a) Schematic of eight ANKLE2 orthologs cloned into expression vectors for characterization in HEK293T cells. (b) ANKLE2 orthologs were transfected into HEK293T cells and purified by affinity-purification (FLAG-AP). Interaction with other host proteins was determined by western blot. EV = empty vector transfection control. (c) HEK293T Cells were transfected, and lysate was split into three fractions. Relative abundance of ANKLE2 orthologs in each fraction was determined by western blot. GAPDH was used as a cytosolic marker, SERCA2 as a membrane organelle marker, and Lamin A/C as a nuclear marker. (d) HeLa cells were transfected, immunostained for ANKLE2 (yellow) and the ER-marker SERCA2 (magenta). Cells were imaged by confocal microscopy at 63X. (e) Pearson’s correlation was used to quantify colocalization between ANKLE2 and the ER. Grey dots indicate individual cells. One-way ANOVA with Dunnett’s multiple comparisons test (compared to human ANKLE2), * p < 0.05, **** p < 0.0001.

To test our hypothesis that presence of a TM or interaction with VAPA mediates ER-localization, we performed subcellular fractionation in HEK293T cells (Figure 3c). We found that all orthologs were primarily present in the membrane-bound organelle fraction representing the ER. Only *Hydra* showed any detection in the cytosolic fraction, which is consistent with the lack of a TM or VAPA interaction. We also observed some orthologs in the nuclear/nuclear lamina fraction, corresponding to the known localization of ANKLE2 to the inner nuclear membrane. This pattern was reproducible in HeLa cells when analyzed by confocal microscopy. All orthologs except *Hydra* demonstrated strong colocalization with the ER-marker SERCA2 as measured by Pearson’s correlation coefficients (Figures. 3d-e, S1). Together, these results show that even very evolutionarily distant ANKLE2-orthologs are ER localized, until Cnidaria where it appears to be diffuse and untethered.

Next, we tested if these orthologs were conserved enough to maintain interaction with human PP2A subunits (Figure 3b), as these protein interactions are amongst the most well-studied for ANKLE2 (Asencio et al. 2012; Snyers et al. 2018; Li et al. 2024, 2025; Saeng-chuto et al. 2025). Human ANKLE2 interacts with the PP2A complex subunits PPP2R1A, PPP2CB, and PPP2R2D, with evidence showing these interactions are mediated by the CD (Asencio et al. 2012; Saeng-chuto et al. 2025). We observed that the closely related mouse and zebrafish orthologs maintained clear interactions with the scaffolding subunit PPP2R1A and catalytic subunit PPP2CB. We chose to include acorn worm amongst these due to its abnormally short ANK. In fruit flies, the ANK cooperates with the CD to mediate interaction with PP2A subunits (Li et al. 2025). In line with this, the acorn worm ortholog had dramatically reduced interaction with PPP2CB compared to others. The horseshoe crab ortholog was the sole ortholog identified without a CD and does not interact with PP2A, supporting the accepted model that the CD is required for the PP2A interaction. Interestingly, the ortholog from roundworm did not interact with human PP2A proteins, but the even more distant *Hydra* ortholog did. This raises interesting questions about how variable protein architecture between these orthologs might impact their interactions.

### 2.3 The LEM domain arose early in ANKLE2 evolution

While sequence is a traditional tool for analyzing genetic similarity and evolution, we wanted to consider the similarity between protein structures. However, protein structure predictions can include disordered peptide chains with structured domains in different regions of the prediction boundary. These disordered regions make rigid protein alignments challenging. To alleviate this, we used FATCAT (Li et al. 2020) followed by US-align (Zhang et al. 2022, 2025) (FATCATèUS-align) to perform flexible pairwise alignments of our AlphaFold predicted structures and generate a TM-score for each pair (Table S1, see Methods, Section 4.8). TM-scores ≥ 0.5 indicate two structures sharing highly similar overall topologies. Analysis of whole ANKLE2 models revealed modest structural similarity, although it was not convincing or consistent enough to draw any major conclusions (Figure S2). We hypothesized that a large fraction of this variability arose from the numerous disordered regions present throughout ANKLE2 (Figure 2, black lines). To sharpen our analysis moving forward, we considered only smaller portions of the proteins at a time, representing individual domains or structured regions.

First, we evaluated the LEM domain of ANKLE2, which mediates binding with BAF and ANKLE2’s partial localization to the nuclear lamina (Cai et al. 2001). The ANKLE2 LEM domain is composed of three simple alpha (α)-helices, and AlphaFold predictions of non-model ANKLE2 orthologs recapitulate this pattern with high-confidence (Figure 4b). Amino acid conservation of this region between orthologs was considerably higher than those of the entire ANKLE2 coding sequence, especially amongst more advanced Chordates (Figure 4a). Overall sequence identity dramatically dropped beginning with the jawless hagfish, which represents more primitive craniates.

**Figure 4:**
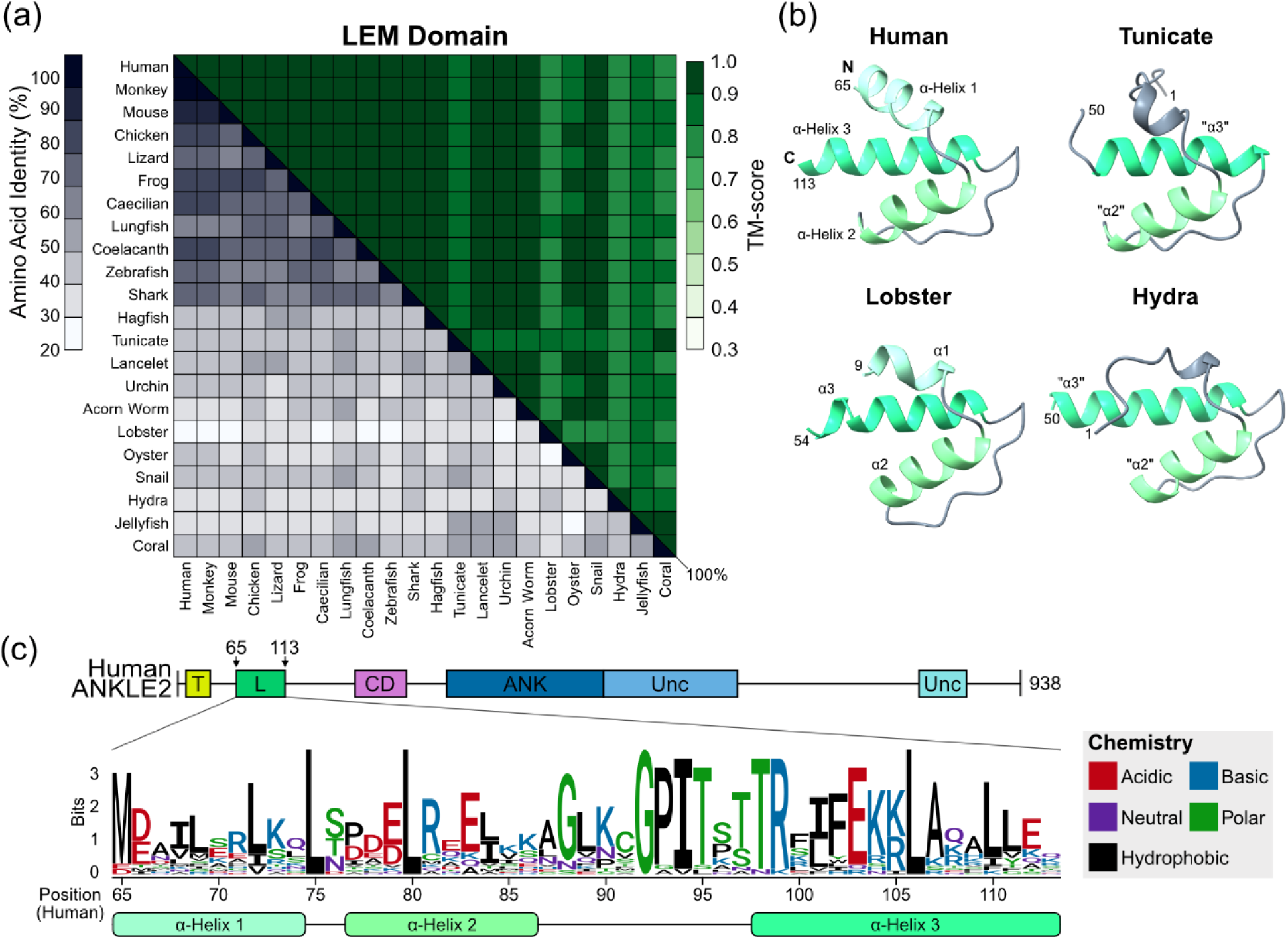
Sequence and structural alignments of the ANKLE2 LEM domain. (a) Sequence-structure matrix of the LEM domain from all relevant orthologs. MSA was computed using Clustal Omega (Madeira) and structural similarity was determined using US-align (REF). (b) Visualization of LEM domains from human, representative of “normal” LEM orientation, compared to “LEM-like” structures predicted in other organisms. (c) Logo plot showing amino acid conservation across the LEM domain.

Despite this divergence in sequence, LEM domain structure predictions were highly similar. US-align structural similarity confirmed that nearly all these LEM domains were identical (Figure 4a). To our surprise, we could identify primitive forms of the LEM domain as far back as Cnidaria, although all three of these only had two of the three helices observed in other LEM domains (Figure 4b). This pattern was shared by tunicate and lobster LEMs which had similar truncated or disordered portions in place of the first α-helix (Figure 4a-b). Logo analysis of these LEM domains revealed that certain amino acids had remarkably high conservation. This conservation was not constrained to only the helices, with much of the highest conservation existing in the unstructured region between helix 2 and 3 (Figure 4c).

While the LEM is crucial for interaction between human ANKLE2 and BAF (Saeng-chuto et al. 2025), multiple other studies dissecting the relationship between ANKLE2, BAF, and cell division were done in fruit flies or roundworms (Asencio et al. 2012; Snyers et al. 2018; Link et al. 2019; Li et al. 2025), which lack any sign of the LEM domain. This suggests that the role of LEM in this interaction is more complex than imagined. Based on the data we collected, we consider the most parsimonious explanation to be that the LEM domain in ANKLE2 arose at least four times, in Cnidaria, Deuterostomia, Mollusca, and *Malacostraca*.

### 2.4 The ankyrin repeat domain is highly conserved among ANKLE2 orthologs but divergent from other ankyrin proteins

The ankyrin repeat domain (ANK) of ANKLE2 spans ∼170 amino acids and contains three distinct ankyrin repeats. Individual ankyrin repeats are composed of two α-helices, an inner and outer helix, and an extended loop called the finger (Mosavi et al., 2004). Vertebrate ANKLE2 orthologs share considerable amino acid identity ranging from 73-88%. Lower in our evolutionary tree, this identity drops dramatically to the range of ∼30-50% between orthologs (Figure 5a). Thus, it was surprising to find that when we compared ANK structures, we found them all to be nearly perfect matches with TM-scores ranging between 0.887-0.997 for all pairs, with the notable exception of the ANK from the hemichordate acorn worm (Figure 5a). The acorn worm ANK region is missing 64 amino acids (relative to human) which correspond to the entire first ankyrin repeat (Figure 5b). We speculate that this missing region impacts this ortholog’s capability to interact with human PP2A complex subunits (Figure 3b), although whether this impacts function within acorn worms themselves is unclear.

**Figure 5:**
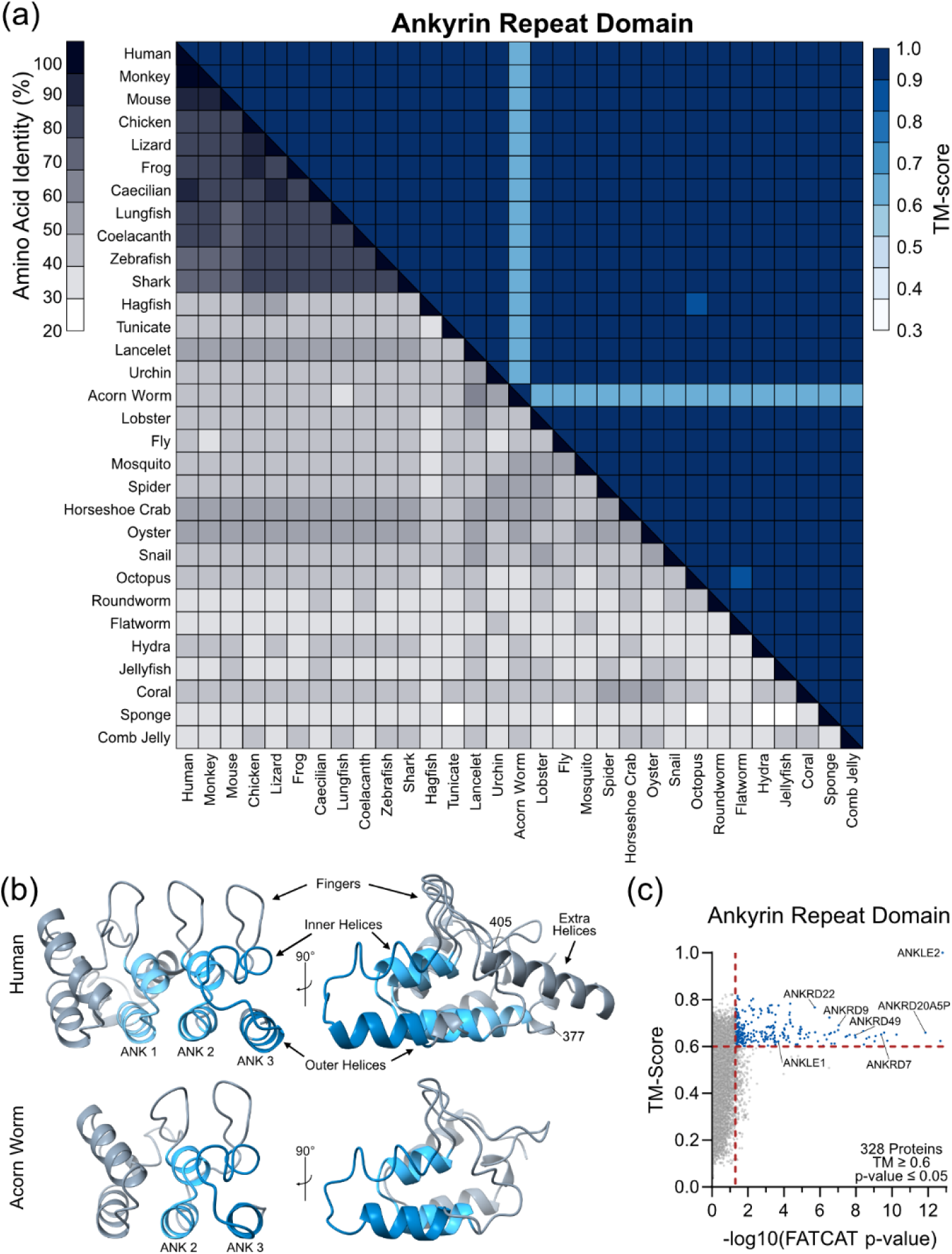
Sequence and structural alignments of the ANKLE2 ankyrin repeat domain. (a) Sequence-structure matrix of the ankyrin repeat domain from all ANKLE2 orthologs. (b) Representative visualization of human ankyrin repeat domain, highlighting helix and finger orientation. Acorn worm is the only abnormal ankyrin repeat domain observed, which clearly lacks the “first” ankyrin repeat. (c) Flexible alignments confirm similarity between ANKLE2 and other known ankyrin repeat domain-containing proteins.

In our investigation of the ANK region, we observed additional high-confidence α-helices (amino acids 377-405 in humans) starting from the outer helix of the first repeat and ending at the start of the second repeat finger (Figure 5b). This addition between ankyrin repeats was surprising. We next investigated if any other proteins had similar insertions within ankyrin repeats, which might provide insight into uniqueness, function, and evolutionary trajectory. We did so by using FATCATèUS-align to compare the ANKLE2 ankyrin region (amino acids 299-473) to the entire human proteome and identify similar structures. This yielded 328 unique proteins above our US-align TM-score and FATCAT p-value cutoffs (Figure 5c). Of these, 87 are named as ankyrin repeat domain proteins, highlighting the accuracy of this approach (Table S1). Despite this, no other proteins had a TM-score greater than 0.82 when compared to ANKLE2, supporting its uniqueness even amongst ankyrin repeat proteins.

### 2.5 Broad-scale structural similarity searches reveal connection between ANKLE2 uncharacterized regions and GIY-YIG nucleases

We next explored if other structured domains in ANKLE2 could be studied in a similar manner using protein structure alignments, potentially revealing their unique structures or shared functions with other well characterized proteins. We used FATCATèUS-align to compare each ANKLE2 domain against the human proteome (Table S1, Figure S3a). This extensive analysis revealed a wide array of human proteins that had structural overlap with portions of ANKLE2. As expected, known LEM and ankyrin repeat proteins were among the top hits when we searched with those ANKLE2 domains (Figure S3b and Figure 5c), supporting the accuracy of this analysis.

One of the largest unexplored aspects of ANKLE2 biology is the two high-confidence structures at the center (“Unc-1”) and C-terminus (“Unc-2”) of the protein. Like other ANKLE2 structured domains, both regions have remarkable structural conservation across animals (Figure S4, S7). Despite these two domains being spatially separated in amino acid sequence, in all structural predictions we evaluated they were closely associated with each other, suggesting they may act cooperatively (Figure 6a). Interestingly, the comb jelly, the evolutionarily “most distant” animal ortholog we analyzed relative to humans (Schultz et al. 2023), has these two domains sequentially contiguous without impacting the predicted structure (Figure 2, 6b). This suggests that these two domains may have initially evolved as a single sequential element and split into two separate domains early in animal evolution.

**Figure 6:**
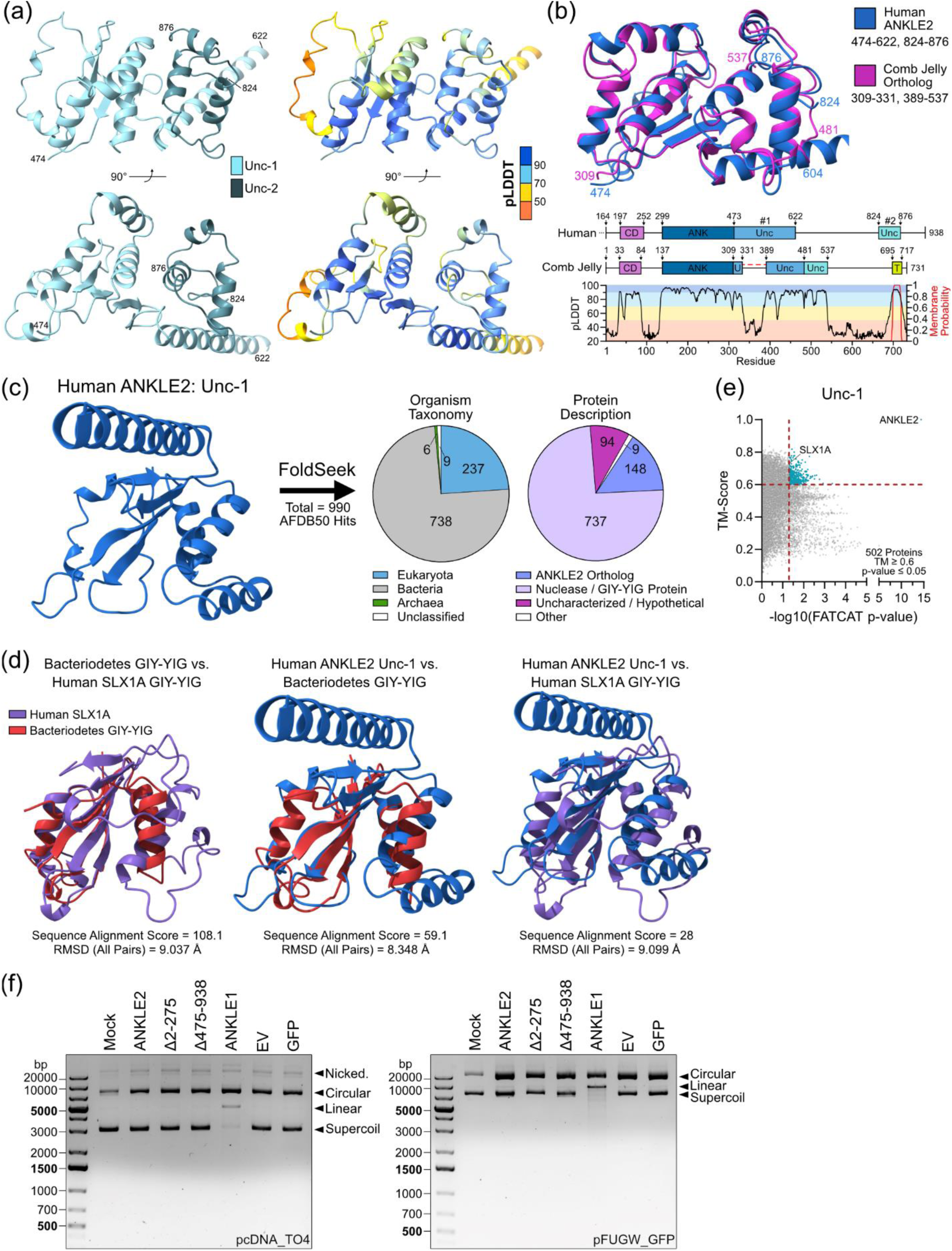
Uncharacterized domain #1 resembles a GIY-YIG nuclease domain but is not enzymatic. (a) AlphaFold representation of human ANKLE2 uncharacterized domains. (b) Structural alignment and protein maps of human and comb jelly ANKLE2. (c) Foldseek results from human ANKLE2 uncharacterized domain #1. (d) Structural alignments of bacterial GIY-YIG nuclease, human SLX1A GIY-YIG domain, and ANKLE2 uncharacterized domain #1. (e) US-align flexible alignment of human ANKLE2 uncharacterized domain #1 confirms similarity with SLX1A. (f) Endonuclease assay in which purified protein was incubated with plasmid DNA to evaluate potential GIY-YIG domain activity of ANKLE2.

To identify its closest similarity outside of humans, we next used Foldseek (van Kempen et al. 2024), an efficient but rigid structure alignment tool, to search across the available AlphaFold proteome without species restriction. Surprisingly, we found consistently high structural overlap between Unc-1 and proteins from a wide array of prokaryotic organisms (Figure 6c). Further, most of these hits were GIY-YIG domain-containing nuclease proteins. These proteins are involved in multiple processes including DNA repair, recombination, and foreign DNA restriction (Dunin-Horkawicz et al. 2006). Interestingly, only two human proteins have annotated GIY-YIG domains, the endonucleases SLX1A and ANKLE1. Aligning the GIY-YIG domain from one of the *Bacteroidetes bacterium* endonuclease proteins with human SLX1A highlights the conservation of this domain across the tree of life (Figure 6d, left panel). Aligning these regions with ANKLE2 Unc-1 revealed modest overlap, sparking our curiosity that this structure may be functionally related to a GIY-YIG domain (Figure 6d, center and right panel).

To explore the similarity between ANKLE2 Unc-1 and human GIY-YIG nucleases we performed flexible alignments of Unc-1 across the human proteome using FATCATèUS-align. We found SLX1A amongst the highest scores (Figure 6e). However, the gap between ANKLE2 and any other protein was vast, suggesting Unc-1 is a structure highly unique to ANKLE2 within the human proteome. Sequence alignments of metazoan SLX1A (Figure S5a) and ANKLE1 (Figure S5b) clearly indicate the highly conserved “GIY” and “YIG” motifs, as well as conserved arginine and glutamic acid residues that are crucial for function (Dunin-Horkawicz et al. 2006). While certain positions of ANKLE2 Unc-1 are highly conserved, we could not identify clear GIY or YIG motifs (Figure S5c), suggesting that while the GIY-YIG nuclease domain is the closest relative to Unc-1, it might not share an overlapping function.

We next tested if ANKLE2 lacks non-specific endonuclease function despite its structural similarity. We expressed and purified ANKLE1, ANKLE2, and two ANKLE2 truncations in HEK293T cells. The first truncation, Δ2-275, lacks the TM, LEM, and CD domains which we thought might be important for DNA recognition based on the known role of LEM in ANKLE1 endonuclease activity (Brachner et al. 2012), while the second truncation, Δ475-938, lacks Unc-1 and Unc-2 (Figure S6). When ANKLE1 was combined with plasmid DNA, we clearly observed endonuclease activity through the production of linearized DNA, as had been previously demonstrated (Brachner et al. 2012) (Figure 6f). However, we did not detect any meaningful changes in DNA migration with ANKLE2. These data, along with the lack of an obvious GIY or YIG motif, suggest that Unc-1 of ANKLE2 does not function as an enzymatic endonuclease domain despite its structural similarity to this domain in other nucleases (Figure 6f).

When analyzing Unc-2 alignments, Foldseek hits were more limited than Unc-1 and primarily eukaryotic proteins (Figure S8). Besides other ANKLE2 orthologs, the next most abundant hits were elongation factor 1-beta (EF-1ß) type proteins. This finding was supported by our flexible FATCATèUS-align alignments of this small domain against the human proteome, which revealed human EF-1ß (EEF1B2) as the strongest hit amongst at least 14 glutathione S-transferase (GST) proteins (Figure S8). It remains unclear whether ANKLE2 has any functional similarity with these proteins.

### 2.6 Key sequence and structural differences in ANKLE2 and RNASEH1 CD distinguish nucleic acid binding ability

The CD of ANKLE2 is a peculiar region lying between the LEM and ANK domains. We identified this region in nearly every ANKLE2 ortholog we examined, with the only exception being horseshoe crab which starts at the ANK (Figure 2). This region is clearly crucial for ANKLE2 function in humans, as multiple mutations in this region are known to cause congenital microcephaly (Fishburn et al. 2024). However, the function of this region in ANKLE2 has only been minimally explored. CD mediates binding with PP2A subunits (Asencio et al. 2012; Saeng-chuto et al. 2025), but this is unlikely to be its only function. Logo analysis of all ANKLE2 orthologs, revealed high sequence conservation throughout the region. However, the space between the first beta (ß)-strand (ß1) and the first α-helix (α1) was variable in length and sequence (Figure 7a), which we designated as the “variable linker”. As with other domains, structural alignments of the CD across ANKLE2 orthologs were consistently high scoring (Figure 7b). This region is composed of two α-helices (α1 and α2) which sandwich two ß-strands (ß1 and ß2) (Figure 7c). The variable linker region loops around from the end of ß1 to the beginning of α1.

**Figure 7:**
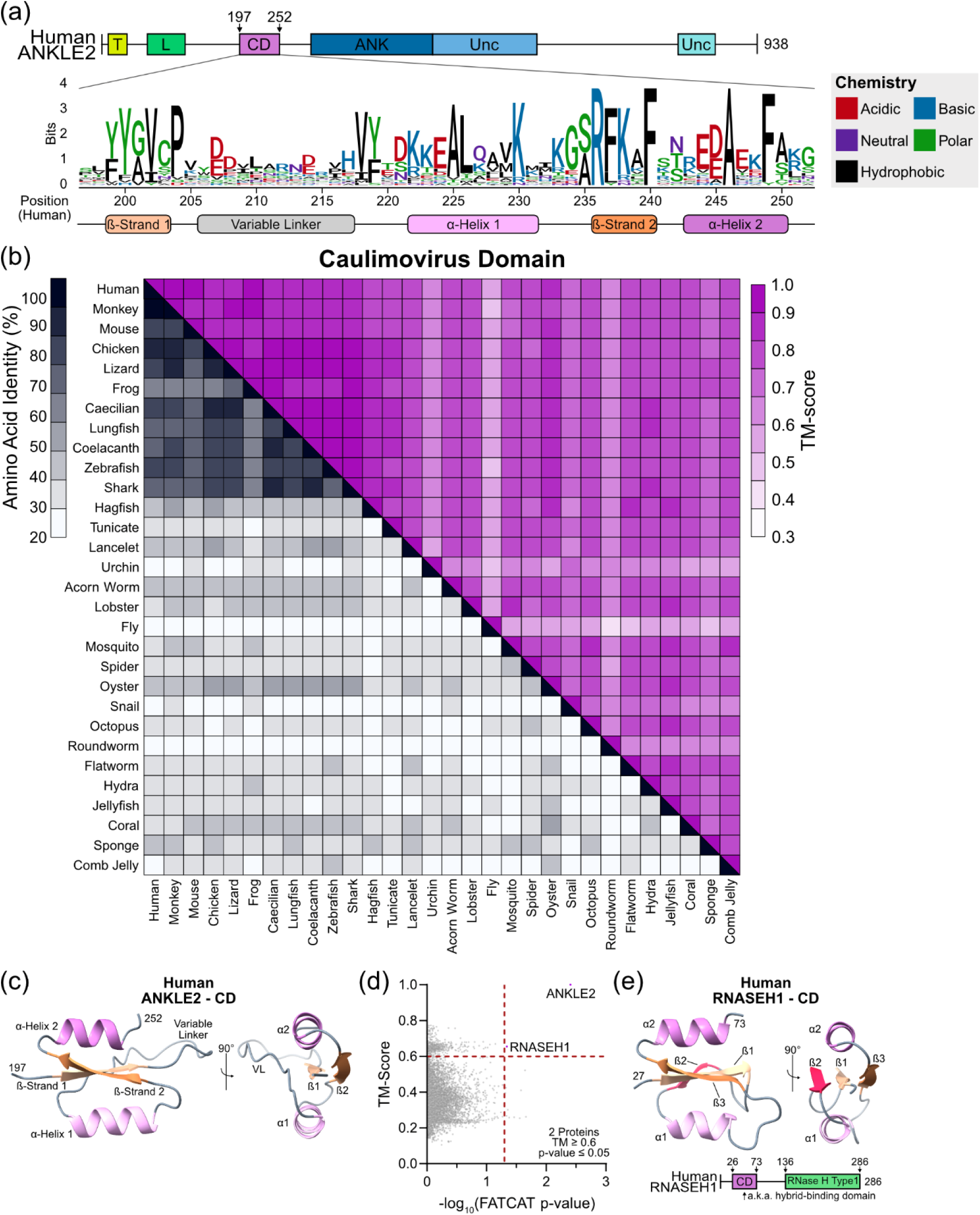
Structural alignments of ANKLE2-CD reveal similarity with RNASEH1 hybrid-binding domain. (a) Logo plot showing amino acid conservation across the CD. (b) Visualization of human ANKLE2-CD. (c) Sequence-structure matrix of the CD from all ANKLE2 orthologs. (d) US-align flexible alignments of human CD reveal RNASEH1 as the only other high-scoring protein. (e) Schematic and visualization of CD from human RNASEH1.

Our structural alignment of CD revealed only one human protein with similar structure, the ribonuclease RNASEH1 (Figure 7d). InterPro and PFAM databases also confirmed that CD is unique to ANKLE2 and RNASEH1 in the human proteome. However, RNASEH1-CD contains a third ß-strand in place of the variable linker we observed in ANKLE2 (Figure 7e). RNASEH1-CD is known to mediate binding to RNA:DNA hybrids, also known as R-loops, which are produced during transcription. These are then degraded by a separate enzymatic domain of RNASEH1 (Nowotny et al. 2008). RNase H proteins are ubiquitous and found throughout the entire tree of life, although it remains unclear whether the CD within RNASEH1 arose from viruses via horizontal gene transfer or was acquired from an even older ancestor. However, ANKLE2 is limited only to animals, and its CD is slightly different from RNASEH1. This suggests that the acquisition of CD within ANKLE2 to be a separate and more recent evolutionary event.

We next explored whether the CD of RNASEH1 and ANKLE2 have similar functions. RNASEH1-CD binds RNA:DNA hybrids and dsDNA with lower affinity (Nowotny et al. 2008). Because the binding interface of RNASEH1 with RNA:DNA hybrids is well characterized, we performed sequence alignments of several ANKLE2-CD and RNASEH1 sequences to look for a similar binding motif. Many of the residues in the hydrophobic core and those important for DNA/RNA interaction are conserved across both proteins. However, the crucial RNA-binding loop of RNASEH1 (residues DRPFA) is absent in ANKLE2-CD. Instead, this region of ANKLE2-CD contains residues of drastically different biochemistry (KMIKG) (Figure 8a). Thus, we hypothesized that despite structural similarities, ANKLE2 would not bind nucleic acids as RNASEH1 does.

**Figure 8:**
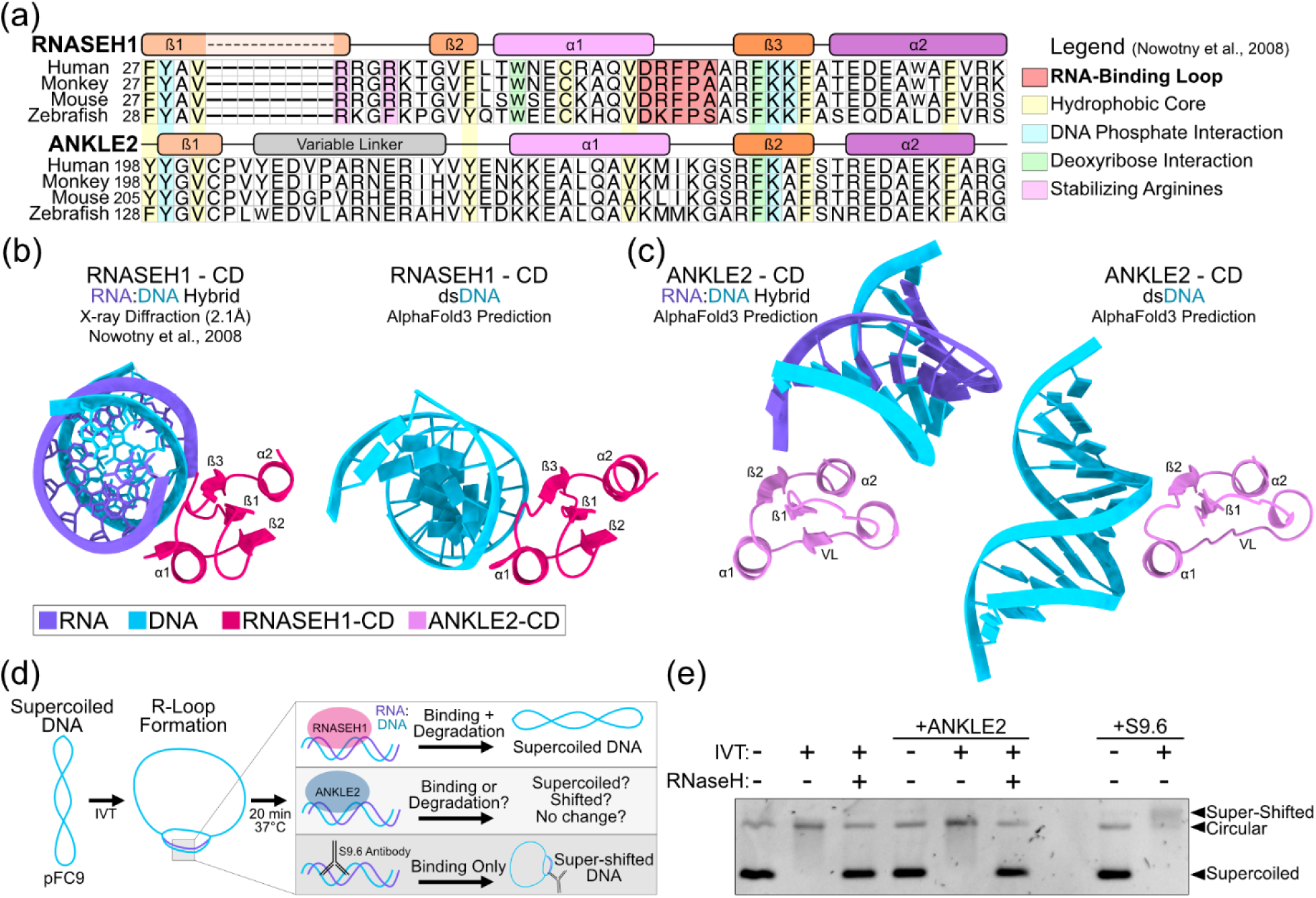
ANKLE2 does not bind RNA:DNA hybrids. (a) MUSCLE amino acid sequence alignment of CD from four species of RNASEH1 and ANKLE2. Colored using designations from Nowotny et al., 2008, highlighting a lack of RNA-binding loop in ANKLE2. (b-c) AlphaFold3 interaction predictions with either RNA:DNA hybrid (GACACCUGAUUC) or dsDNA (GACACCTGATTC) compared to experimentally derived structure of RNASEH1 interacting with hybrid (Nowotny et al. 2008). RNA is shown in purple and DNA in cyan. VL = variable linker, α1/2 = alpha-helix 1 and 2, ß1/2 = beta-strand 1 and 2. (d) Visual schematic RNA:DNA hybrid binding-degradation assay. (e) Hybrid binding assay to assess ability of purified human ANKLE2 to bind or degrade R-loops produced during *in vitro* transcription.

We first tested this hypothesis computationally. AlphaFold3 predictions of RNASEH1-CD with dsDNA produced complexes with the same binding interface as experimentally resolved structures of hybrid binding (Figure 8b) (Nowotny et al. 2008). When we attempted this prediction with ANKLE2-CD, we found that the contact interface between both RNA:DNA hybrid and dsDNA were completely different and likely an artifact. This suggests that despite the surprising structural similarities between ANKLE2-CD and RNASEH1-CD, it does not bind nucleic acids in a similar fashion (Figure 8c).

To test this prediction experimentally, we performed *in vitro* binding assays with dsDNA and RNA:DNA hybrids. First, we tested binding to non-specific dsDNA using electrophoretic mobility shift assay (EMSA), which revealed no convincing evidence of DNA binding (Figure S9). Next, to test binding to RNA:DNA hybrid, we used the plasmid pFC9 and generated stable R-loops using *in vitro* transcription (IVT) with T7 RNA Polymerase (RNAP) (Ginno et al. 2012; Stolz et al. 2019) (Figure 8d). All IVT reactions were done with supercoiled plasmid. R-loop formation was confirmed by a tell-tale loss in electrophoretic mobility upon addition of T7 RNAP. This loss of mobility was restored upon treatment with purified bacterial RNase H, confirming R-loop production and RNase H activity (Figure 8e, lanes 1-3). When we added purified ANKLE2, we did not observe any change in plasmid mobility in either un-transcribed or transcribed samples. These data indicate that in contrast to RNase H, ANKLE2 was not acting as an R-loop resolvase under these conditions (Figure 8e, lanes 4-6). Though unlikely based on the AlphaFold predictions, ANKLE2 may bind R-loops but be unable to degrade them without a functional nuclease domain. To test this specifically, we compared our ANKLE2 treatment to samples incubated with the antibody S9.6 (Boguslawski et al. 1986) which stably binds R-loops and produces super-shifted bands (Figure 8e, lanes 7-8). We did not observe any super-shifted bands when un-transcribed or transcribed samples were incubated with ANKLE2. This, in combination with comparison to RNase H treatment of R-loop containing plasmids, suggests that it does not bind or degrade RNA:DNA hybrid nucleic acids.

### 2.7 Depletion of ANKLE2 results in reduced DNA damage and increased micronuclei formation

Depletion of RNASEH1 results in DNA damage (Amon and Koshland 2016; Ohle et al. 2016p.201). Despite our evidence that ANKLE2-CD did not directly bind nucleic acids, we sought to investigate if ANKLE2-CD confers similar roles in DNA damage responses. We used ANKLE2 knockout (KO) Huh7 cells we previously generated (Fishburn et al. 2025) and treated them with etoposide. Etoposide inhibits topoisomerase II, leading to the accumulation of DNA damage in the form of double-strand breaks (DSBs) (Montecucco et al. 2015). We then visualized and quantified the number of DSBs using γ-H2AX, an antibody that targets the phosphorylated variant of the H2A histone protein present at DSBs (Figure 9a). We found that ANKLE2 KO cells were less sensitive to etoposide, with fewer γ-H2AX foci at two concentrations of etoposide and even in our DMSO control (Figure 9b), suggesting that either ANKLE2 KO cells are more resistant to DNA damage or that ANKLE2 may repress the response to DNA damage. It is unclear if this is an indirect mechanism or if ANKLE2 is directly involved in coordinating the phosphorylation of histone proteins in response to DSBs. VRK1 phosphorylates H2AX at serine 139 in response to DNA damage (Salzano et al. 2015), providing a possible explanation for this result.

**Figure 9:**
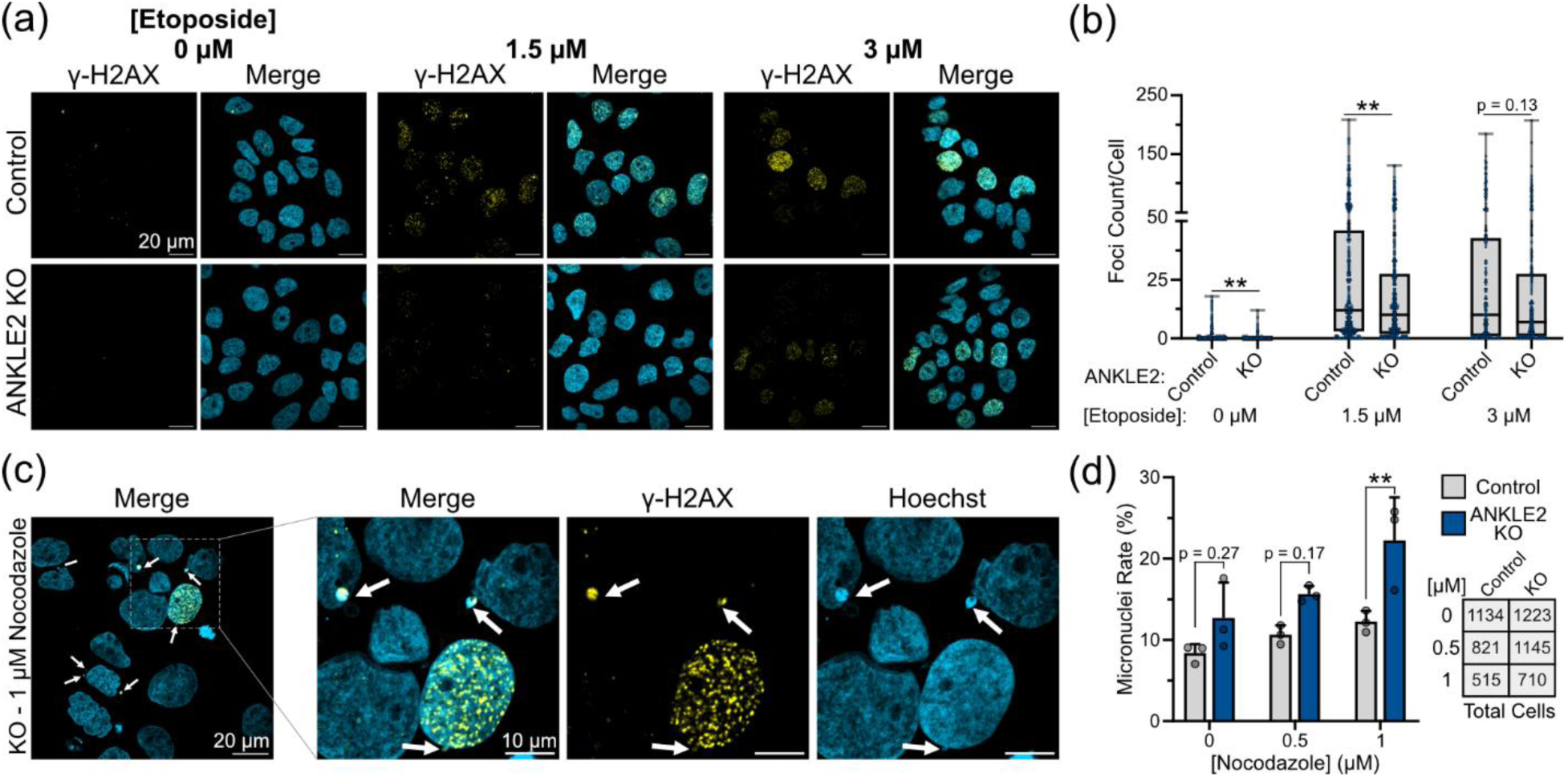
Depletion of ANKLE2 confers DNA damage resistance but sensitizes cells to micronuclei formation. (a) Representative confocal images of Huh7 control or ANKLE2 knockout (KO) cells that were treated with etoposide for 1 hour prior to fixation and immunostaining. (b) DNA damage foci were counted using an automated imaging pipeline. Blue dots indicate individual cells. (c) Representative image of Huh7 ANKLE2 KO cells treated with nocodazole for 24 hours prior to fixation and immunostaining. (d) Micronuclei were counted manually from images taken at 40X or 63X. Grey dots indicate biological replicate experiments where 10 images were taken from each experiment and micronuclei rate (cells with at least one micronuclei / total cells) was calculated.

During this experiment we also observed that many cells had micronuclei (MN), small DNA-containing structures present outside the primary nucleus. MN can arise due to defects in spindle assembly, nuclear envelope dynamics, or chromosome segregation (Krupina et al. 2021), and are common in cancers and cancer cell lines such as Huh7. In our etoposide treatment experiment, we observed that ANKLE2 KO cells consistently had more MN than control cells (Figure S10). To confirm this phenotype, we treated the same cells with nocodazole, a microtubule polymerization inhibitor, known to induce MN (Zhang et al. 2015). Again, we observed that ANKLE2 KO cells consistently had more MN than control cells (Figure 9c-d). Together, we conclude that ANKLE2 is involved in the regulation of DNA damage and MN formation.

## 3 DISCUSSION

ANKLE2 is a protein of emerging importance in multiple fields of study (Fishburn et al. 2024) and as its study expands across multiple model organism systems (Asencio et al. 2012; Link et al. 2019; Apridita Sebastian et al. 2022; Li et al. 2025), so does the necessity to understand how ANKLE2 has evolved. In this work, we present a detailed description of ANKLE2 orthologs from over 30 species to draw conclusions about its evolutionary history. By expressing some of these orthologs *in vitro*, we were able to characterize their subcellular localization and the conservation of key protein interactions. To investigate the conservation of each domain of ANKLE2 we compared sequence identity and structural similarity using computational approaches. By expanding our structural conservation analysis to all proteins, we identified similarities with RNASEH1 and GIY-YIG nucleases. The structure-function relationship prompted us to further probe if these domains conferred similar biochemical properties to ANKLE2. Despite the structural similarities, important motifs critical to function were absent. Nonetheless, we believe this structure-forward strategy provides valuable opportunities in uncovering new biological phenomena, such as ANKLE2’s regulation of DNA damage. Use of structural prediction as a tool in phylogenetic analysis is growing more common as the technology becomes more widespread and accessible (Kim et al. 2025; Mishra et al. 2025; Hunker and Bleichert 2026).

One of our most promising conclusions is that the TM in ANKLE2 emerged recently during the evolution of Chordates. Based on our results, we speculate that ANKLE2 evolved a membrane anchoring mechanism through interaction with tethering proteins like VAPA in an early common ancestor of protostomes and deuterostomes. During chordate development, the N-terminal TM emerged as an alternative mechanism to VAPA interaction. Since the subcellular localization of the ANKLE2 is highly important in regulating cell division and brain development, a self-contained TM may have proved advantageous as neurodevelopmental complexity evolved. It is worth noting that we did identify predicted TM domains in both the comb jelly and octopus orthologs, suggesting at least two additional independent evolutionary events of an ANKLE2 TM. We predict these domains to facilitate ANKLE2 membrane anchoring, though the curious C-terminal position of the TM in comb jelly suggests an alternative membrane insertion mechanism to maintain cytosolic/luminal topology.

The evolution of ANKLE2’s LEM domain is perplexing, despite it being much of the protein name and crucial for its canonical interaction with BAF. Nearly all the biochemical and cellular functions of ANKLE2 attributed to cell division, nuclear envelope dynamics, and coordination of BAF phosphorylation were performed in fruit flies or roundworms (Asencio et al. 2012; Snyers et al. 2018; Link et al. 2019; Li et al. 2025), both of which completely lack any semblance of a LEM domain. One possible explanation is that these orthologs mediate interaction with BAF through another domain, and LEM evolved later to strengthen the protein interaction or improve localization to the nuclear lamina. However, this is confounded by the observation of LEM-like structures in Cnidaria and *Malacostraca* in the same N-terminal location (Figure 2).

It is tempting to attribute the unexplained function of these ANKLE2 orthologs to the uncharacterized domains, which are present and structurally conserved throughout the animals we evaluated. Our analysis revealed similarities between Unc-1 and GIY-YIG nucleases (Figure 6). Given that ANKLE1 also functions as a nuclease, we were especially hopeful to identify a similar novel function in ANKLE2. However, the classic GIY-YIG motifs are not present, resulting in no clear enzymatic function. We speculate this domain serves in supporting ANKLE2’s scaffolding function, but further study is required to interrogate the true nature of this domain. The importance of this domain is underscored by a recent clinical case in which compound heterozygous mutations at R536C/G566R resulted in *ANKLE2-*related congenital microcephaly (Barsh et al. 2025).

Unc-2 remains equally mysterious; however, its similarity to EF-1ß and GST-binding proteins is a promising lead. Previous study in fruit flies has attributed the interaction with VAPA to this region (Li et al. 2025). However, the presence of this domain alone does not appear to be sufficient for interaction with VAPA, since human and mouse Unc-2 fail to mediate an interaction with VAPA (Figure 3b). In humans, mutations in both uncharacterized regions are pathogenic and result in neurodevelopmental disease (Thomas et al. 2022; Fishburn et al. 2024), highlighting that novel functions remain to be discovered.

The final goal of our study was to explore ANKLE2-CD. Its unique structure was revealed to only be shared with RNASEH1, where it mediates RNA:DNA binding (Nowotny 2018). Despite the prominent similarity between ANKLE2-CD and RNASEH1-CD, critical RNA-binding residues were absent in ANKLE2-CD (Figure 8c). Consistent with the absence of these critical residues, ANKLE2 did not bind dsDNA or RNA:DNA. Together with our findings with Unc-1 and the lack of a clear GIY or YIG motif, our work highlights that broad structural similarity across a domain does not necessarily indicate a conserved function. Despite this, we were able to show that ANKLE2 regulates aspects of DNA-damage response and MN formation (Figure 9). However, since these functions were revealed in ANKLE2 KO cells, it is possible that some other domain outside of CD is responsible for regulating DNA damage.

It is important to acknowledge the limitations of our study as it pertains to our results and conclusions. At the forefront is our usage of AlphaFold and structural predictions in general, which heavily rely upon existing resolved structures as training data. While model organisms such as roundworms, fruit flies, and humans comprise large swaths of available datasets, niche organisms referenced in this paper lack the same depth of understanding. AlphaFold2 and the recent version 3 offer promising opportunities for computational discovery, but they do not always reflect physical structures and associated biological functions. We have combined modern computational approaches with traditional biochemical assays to mitigate these shortcomings.

Despite the advantages of combining computational and experimental approaches, our experimental approaches have their own limitations. Expression of ANKLE2 orthologs in human cells provides us with the opportunity to evaluate their behavior in a cellular context. This is especially valuable for non-model organisms where facile cell culture models may not be readily available. However, by expressing them in human cells rather than that of the original species, we may lose any specific contributions of ANKLE2-protein interactions that are species-specific. Finally, our ANKLE2 biochemical assays may be complicated by the difficulty in purifying ANKLE2 in a way that maintains its native folding. As discussed, ANKLE2 has intrinsically disordered regions and is membrane-incorporated, both of which increase purification difficulty without compromising stability or function. Ultimately, we identified a purification strategy which yielded protein with minimal degradation (Figure S6) and positive results for ANKLE1 activity (Figure 6), alleviating most of our concern regarding ANKLE2 protein purification.

Together, our work shows that despite variable amino acid sequence conservation, individual ANKLE2 domains are structurally conserved across nearly all animals. This supports a paradigm in which protein structure similarity, rather than sequence identity/similarity alone, is a valuable metric evaluating protein conservation and evolution. However, this is met with an equally worthy disclaimer that, just because proteins share similar structures, does not mean that they necessarily share similar functions. Understanding which biological properties are conferred inherently by structure versus which are conferred by the specific activity of amino acid side chains is crucial in differentiating when the structure-function relationship is conserved and when it is not.

## 4 MATERIALS AND METHODS

### 4.1 Protein sequence collection, structure prediction, and domain annotation

Protein sequences were collected from NCBI and accession numbers of all used sequences are in Table S2. Protein structures were predicted using AlphaFold2 through Google Collab (ColabFold version 1.5.5: AlphaFold2 using MMseqs2) or downloaded from the EMBL-EBI database (https://alphafold.ebi.ac.uk) when available. *Homo sapiens* proteins for the ANKLE2 vs. All searches were downloaded as a bulk set from the AlphaFold EMBL-EBI database. Transmembrane domains were annotated using DeepTMHMM (Hallgren et al. 2022). Other structural domains (ANK, CD, LEM, Unc) were annotated by examining pLDDT values, α-helix, and ß-sheet determinations from AlphaFold predictions along with sequence alignments with other orthologs. Structural predictions were visualized in ChimeraX v1.6.1 (Pettersen et al. 2021). Protein sequences were aligned with multiple sequence alignments using the MUSCLE algorithm (Edgar 2004) in SnapGene (v4.3.11) or ClustalOmega in the EMBL-EBI online tool (Madeira et al. 2024).

### 4.2 Cells

HEK293T, Huh7, and HeLa cell lines were maintained in Dulbecco’s modified Eagle’s medium (DMEM; Gibco, Thermo Fisher) supplemented with 10% fetal bovine serum (FBS; Gibco, Thermo Fisher) at 37°C, 5% CO2. Cells we washed with Dulbecco’s phosphate-buffered saline (D-PBS, Life Technologies) and dissociated with 0.05% trypsin-EDTA (Life Technologies). Cells were tested for Mycoplasma spp. monthly using a detection kit (Invivogen, NC2326021). CRISPR ANKLE2 knockout (KO) cells were generated and used as previously described (Fishburn et al. 2025).

### 4.3 Plasmids

ANKLE2 ortholog sequences were acquired from NCBI and DNA sequences were codon-optimized for expression in human cells. Gene fragments were synthesized by Twist Biosciences and cloned into pcDNA_TO using Gibson Assembly with KpnI and ApaI (NEB). Plasmid assembly was confirmed with whole plasmid sequencing (Plasmidsaurus). ANKLE2 truncation plasmids were generated using a similar strategy. ANKLE1 was amplified from an existing plasmid (unpublished) using PCR (Forward Primer: TAAGCTTGGTACCGAGCTCGGCCGCCATGTGTTC, Reverse Primer: CGGAACCCCCCCCACTCGCTCCACGCGCTTGAATATCTTGT).

### 4.4 Expression and affinity-purification of ANKLE2 orthologs

Plasmids (7 µg) were transfected into ∼5e6 HEK293T cells in 10 cm dishes using 21 µL PolyJet (SignaGen, NC1536117) for 24 hours. Media were then removed from each plate. To dissociate cells, 5 mL of D-PBS supplemented with 10 mM EDTA was added and allowed to incubate for several minutes. Cells were resuspended in 5 mL of D-PBS and transferred to 15 mL conical tubes prior to centrifugation at 94 × g, 4°C for 5 min (Eppendorf centrifuge 5810 R, Rotor S-4–104). Cell pellets were washed with 5 mL D-PBS and centrifugation was repeated. Supernatant was removed, and pellets were then resuspended in 1 mL IP buffer (50 mM Tris base, 150 mM NaCl, 0.5 M EDTA, pH 7.4) with Pierce protease inhibitor tablets (Thermo Scientific, PIA32955) supplemented with 0.5% NP-40 Substitute (Igepal CA-630, Affymetrix). Cells were lysed for 30 min at 4°C, and lysate was then centrifugated at 845 × g, 4°C, for 20 min (Eppendorf centrifuge 5424 R, Rotor FA-45–24-11). A portion of each lysate (60–100 μL) was collected and saved for Western blot analysis. The remaining lysate was added to 40 μL of magnetic FLAG beads (Sigma, M8823) and incubated overnight at 4°C with gentle rotation.

Beads were then washed four times with 1 mL IP buffer with 0.05% NP-40 and once with 1 mL IP buffer without NP-40. Beads were then incubated in 40 μL of 100 ng/mL FLAG peptide (APExBIO) at 211 × g for 1 hour at room temperature (Eppendorf ThermoMixerC). The eluate was then collected. For purification of ANKLE2 and other proteins for endonuclease and nucleic acid binding assays, this process was repeated with an alternative buffer (50 mM Tris, 300 mM KCl, 5 mM MgCl_2_, 1 mM EDTA, 50mM DTT, 15% glycerol, Pierce protease inhibitor, pH 8.0) which was used in place of IP buffer whenever possible.

### 4.5 Subcellular fractionation of ANKLE2 orthologs

HEK293T cells were transfected as previously described and collected. Suspended cells were subjected to centrifugation (500 × g, 4°C, for 10 min) and pellets were resuspended in 500 µL ice-cold D-PBS. After another spin, cell pellets were resuspended in 400 µL ice-cold “Lysis Buffer A” (150 mM NaCl, 50 mM HEPES, 25 µg/mL digitonin [Neta Scientific, RPI-D43065], 1 M glycerol, Pierce protease inhibitor). Samples were incubated at 4°C for 10 minutes with end-over-end rotation prior to centrifugation at 2000 × g, 4°C, for 10 min. This soluble “Cytosolic” fraction was then collected. Pellets were thoroughly resuspended in 400 µL of ice-cold “Lysis Buffer B” (150 mM NaCl, 50 mM HEPES, 1% Ipegal CA-630, 1M glycerol, Pierce protease inhibitor) and incubated at 4°C for 30 minutes, prior to centrifugation at 7000 × g, 4°C, for 10 min. This soluble “Membrane-bound organelle” fraction was then collected.

Pellets were thoroughly resuspended in 400 µL of ice-cold “Lysis Buffer C” (150 mM NaCl, 50 mM HEPES, 0.5% sodium deoxycholate, 0.1% sodium dodecyl sulfate [SDS], 1M glycerol, Pierce protease inhibitor) and incubated for 30 minutes at 4°C with end-over-end rotation. Samples were then sonicated (Branson Analog Sonifier 250) 2 × 20 second pulses, 95% Duty Cycle with a narrow probe, prior to centrifugation at 7800 × g, 4°C, for 10 min. This final “Nuclear” fraction was then collected.

### 4.6 Western blot and silver stain

Protein samples (lysates or affinity-purification [AP] eluates) were resuspended in NuPAGE LDS sample buffer supplemented with tris(2-carboxyethyl)phosphine (TCEP) and boiled at 95°C for 10 min. Samples were run on 4%–20% gradient polyacrylamide gels for ∼1 hour at 150 V. For western blot proteins were transferred to polyvinylidene fluoride (PVDF) membranes (VWR) for 1 hour at 330 mA on ice. Membranes were then blocked in 5% milk solution for 1 hour prior to overnight incubation in primary antibodies (Table S3) at 4°C. Membranes were washed 3× in Tris-buffered saline with Tween 20 (TBS-T) (150 mM NaCl, 20 mM Tris base, 0.1% Tween 20; Thermo Fisher) and incubated with horseradish peroxidase (HRP) conjugated secondary antibodies in 5% milk for 1 hour at room temperature. Membranes were again washed 3× in TBS-T and 1× in Tris-buffered saline (without Tween 20) prior to Pierce ECL activation (Thermo Fisher). Membranes were imaged using Amersham Imager 600 (GE). For silver stain, the gel was prepared and stained per manufacturer’s recommendations (Thermo Fisher, PI24612). Gels were imaged with a BioRad GelDoc Go Imaging System. Images were analyzed using Fiji (Schindelin 2012).

### 4.7 Immunofluorescence and confocal microscopy

HeLa or Huh7 cells cultured on #1.5 coverslips were fixed with 4% paraformaldehyde (Thermo Fisher) for 15 minutes at room temperature. Cells were permeabilized with 0.1% Triton X-100 (Integra) for 10 minutes and blocked with 5% goat serum (Sigma) in PBS-Tween (0.1% Tween 20, Thermo Fisher). Coverslips were incubated with primary antibodies overnight at 4°C. Coverslips were then washed in PBS-Tween and incubated in secondary antibody at room temperature for 1 hour. Nuclei were visualized with Hoechst (1:10000, Invitrogen). Confocal images were acquired using a Zeiss Airyscan LSM980 with Axiocam using a ×63/NA1.40 oil immersion lens. Laser lines at 405, 488, and 543 nm were employed sequentially for each image using optics and detector stock settings in the “Dye List” portion of the FluoView microscope-controlling software. Other microscopy images were captured using a Nikon Ti2 inverted microscope, CFI PLAN APO LAMBDA ×40 CF160 Plan Apochromat Lambda ×40 objective lens, N.A. 0.95, W.D. 0.17–0.25 mm, F.O.V. 25 mm, DIC, correction collar 0.11–0.23 mm, spring loaded, and using Andor Zyla VSC-08688 camera. All antibodies and dilutions are listed in Table S3. Microscopy images were analyzed using ImageJ (Fiji) software (Schindelin et al. 2012). Signal co-localization was quantified using the BIOP-JACoP plugin (Bolte and Cordelières 2006) within Fiji to quantify Pearson’s correlation coefficient (R value) of individual cells.

### 4.8 ANKLE2 structural alignment across organisms using FATCAT and US-align

We used FATCAT 2.0 (https://github.com/GodzikLab/FATCAT-dist, accessed August 2, 2024.) (Li et al) to perform pairwise flexible alignments for each full-length protein or domain of interest for all organisms with ANKLE2, RNASEH1, etc. orthologs. The output aligned structures were used as inputs to US-align (https://aideepmed.com/US-align/, accessed November 12, 2024) (Zhang et al. 2022, 2025). All parameters for both algorithms were left as default. P-values from FATCAT and TM-scores from US-align were used to determine protein similarity. FATCAT provides a flexible alignment but outputs length-dependent RMSD values. TM-score is length-independent and provides a consistent similarity metric across alignments. It is worth noting that US-align provides a new alignment between two structures. However, we observed the difference in RMSD between alignments to be small in most cases (Table S1).

### 4.9 ANKLE2 structure alignment against the human proteome

We used the same pipeline as stated above to compare human ANKLE2 (UniProt Acc. No. Q86XL3) to the entire human proteome. We performed pairwise alignments with both FATCAT and US-align between ANKLE2 and all available predicted *Homo sapiens* protein structures (23,391 in total) from the AlphaFold2 database (v4). When establishing proteins with similar structures, we used P-value and TM-score. Significance values of -log_10_(P-value) < 1.3 (P-value < 0.05) and TM-score > 0.6 were used as thresholds. TM-scores of > 0.45 or ≥ 0.5 are cited as indicating a high level of similarity between two protein structures (Xu and Zhang 2010; Zhang et al. 2022, 2025), however we maintained a more conservative approach in our alignments by establishing a threshold of TM-score = 0.6.

### 4.10 Endonuclease Assay

Endonuclease assays were performed as previously described (Brachner et al., 2012). In short, 3 µL of purified protein was combined with 150 ng of plasmid DNA (pcDNA_TO or pFUGW_GFP) in endonuclease buffer (20 mM HEPES-KOH, 2mM MnCl_2_, 45mM KCl, 50 µg/mL BSA, pH 7.4) for a total volume of 12 µL. Samples were incubated at 37°C for 60 minutes prior to adding 5 µL of termination buffer (14 mM EDTA, 0.1% SDS, 0.1 mg/mL Proteinase K) and incubating at 55°C for 15 minutes. 2 µL of 10% glycerol and 3 µL of TriTrack DNA loading dye (ThermoFisher, R1161) were added and samples were run on 1% agarose gels for 60 minutes at 80 volts. Gels were imaged with a BioRad GelDoc Go Imaging System.

### 4.11 Assessment of DNA damage and micronuclei

Huh7 cells were plated for immunofluorescence on glass coverslips as previously described and grown overnight. Etoposide was added at indicated concentrations for 1 hour prior to fixation and antibody staining. Gamma-H2aX puncta were counted using an automated imaging pipeline. For micronuclei experiments, nocodazole (Sigma, 487928) was added at indicated concentrations for 24 hours. Cells were then fixed and imaged.

### 4.12 Nucleic acid binding assays

To determine binding with dsDNA electron mobility shift assays (EMSA) were performed according to manufacturer recommendations (ThermoFisher, E33075). In brief, a 511 bp portion of pcDNA_TO was PCR amplified (F: TGGGAGTTTGTTTTGGAACCA, R: CAGATGGCTGGCAACTAGAAG) and purified. 20 ng of this purified DNA was combined with 0/1.5/3 µL of purified protein in binding buffer (150 mM KCl, 0.1 mM DTT, 0.1 mM EDTA, 10 mM Tris, pH 7.4) and incubated at 4°C for 30 minutes. After binding, loading dye was added and samples were run on 4-20% gradient polyacrylamide gels without SDS at 100 volts for ∼5 hours on ice. Gels were then stained in SYBR Green solution (1:10000 stain concentrate in TBE [89 mM Tris, 89 mM boric acid, 1 mM EDTA, pH ∼7.4) for 40 minutes at room temperature, washed, and then imaged. Gels were washed and stained in SYPRO Ruby solution (SYPRO Ruby protein stain in ∼824 mM trichloroacetic acid) overnight at room temperature. Gels were washed and de-stained (10% methanol, 7% acetic acid) for ∼3.5 hours at room temperature, washed, and imaged with a BioRad GelDoc EZ Imager.

To determine binding with RNA:DNA hybrid *in vitro* transcription (IVT) assays were performed using the R-loop forming pFC9 plasmid (Stolz et al. 2019) as described in (Ginno et al. 2012) at a concentration of 20 ng/µL. T7 RNA polymerase (2.5 U/mL, NEB) was added to transcribed samples in the buffer provided by the polymerase manufacturer supplemented with rNTP mix solution (NEB, N0466S) added to a final concentration of 0.5 mM and DTT (NEB, B1222A) to a final concentration of 10 mM. Reactions were incubated at 37°C for 20 minutes and then were terminated by the addition of EDTA to a final concentration of 35 mM. Products were precipitated via phenol-chloroform EtOH precipitation and eluted into RNase H reaction buffer (NEB, B0297SVIAL). Samples were treated with RNase A (33 ng/mL) for 30 minutes at 37°C to degrade excess free RNA. To confirm the presence of R-loops, a portion of the transcribed sample was treated with RNase H (NEB M0297S, 40 ng DNA/1U) for 30 minutes at 37°C to degrade RNA:DNA hybrids. To evaluate ANKLE2 activity on RNA:DNA hybrids, 1 µL of purified ANKLE2 protein was added to 300 ng substrate in a total volume of 30 µL of 1X RNase H binding buffer for 30 minutes at 37°C. As a positive control for RNA:DNA hybrid binding transcribed and un-transcribed pFC9 were incubated with the S9.6 antibody according to (Sanz et al. 2021p.20) for 20 minutes at room temperature. All samples were then loaded and ran on a 0.8% 1X TBE agarose gel at 60 volts for two hours. The gel was then stained with ethidium bromide in 1X TBE buffer for 30 minutes and destained in 1X TBE buffer overnight and imaged with a BioRad GelDoc EZ Imager.

## AUTHOR CONTRIBUTIONS

**Adam T Fishburn**: Conceptualization; formal analysis; investigation; writing – original draft preparation; supervision; visualization; writing – review & editing. **Cole J Florio**: Conceptualization; investigation. **Chase L S Skawinski**: Data curation; formal analysis; investigation; software; writing – original draft preparation. **Sydney S Becker**: Investigation. **Ethan Holleman**: Investigation. **Avery E Robertson**: Investigation, validation. **Rees Sitchon**: Investigation. **Frédéric Chédin**: Resources. **Priya S Shah**: Funding acquisition; supervision; writing – review & editing.

## ACKNOWLEDGEMENTS

This work was supported by funding to P.S.S. provided by the University of California, Davis, and the National Institutes of Health (NIH) (R01/R56AI170857). The Zeiss 980 confocal microscope used in this study was purchased using NIH Shared Instrumentation Grant S10OD026702. We thank the MCB Light Microscopy Imaging Facility, which is a UC Davis Campus Core Research Facility, for the use of this microscope. We thank all our collaborators in the ANKLE2 field for their continued support and shared interest, including Nicole Link and Matthew Evans who helped spark the idea for this work. We thank members of the Shah lab for their constant encouragement and helpful feedback, especially Shruthi Garimella who also helped with AlphaFold3 predictions. We additionally thank members of the UC Davis community including Shawn Christensen for early-stage conception, Jacqueline Barlow for providing reagents and helpful discussion regarding RNase H1, and Celina Juliano and Jasmine Mah for assistance with phylogenetic trees and evolutionary analysis, and Arthur Charles-Orszag for assistance with protein purification and EMSA.

## DATA AVAILABILITY STATEMENT

The data and computational tools used in this study are available upon request.

## Supplementary Material

**Figure S1:**
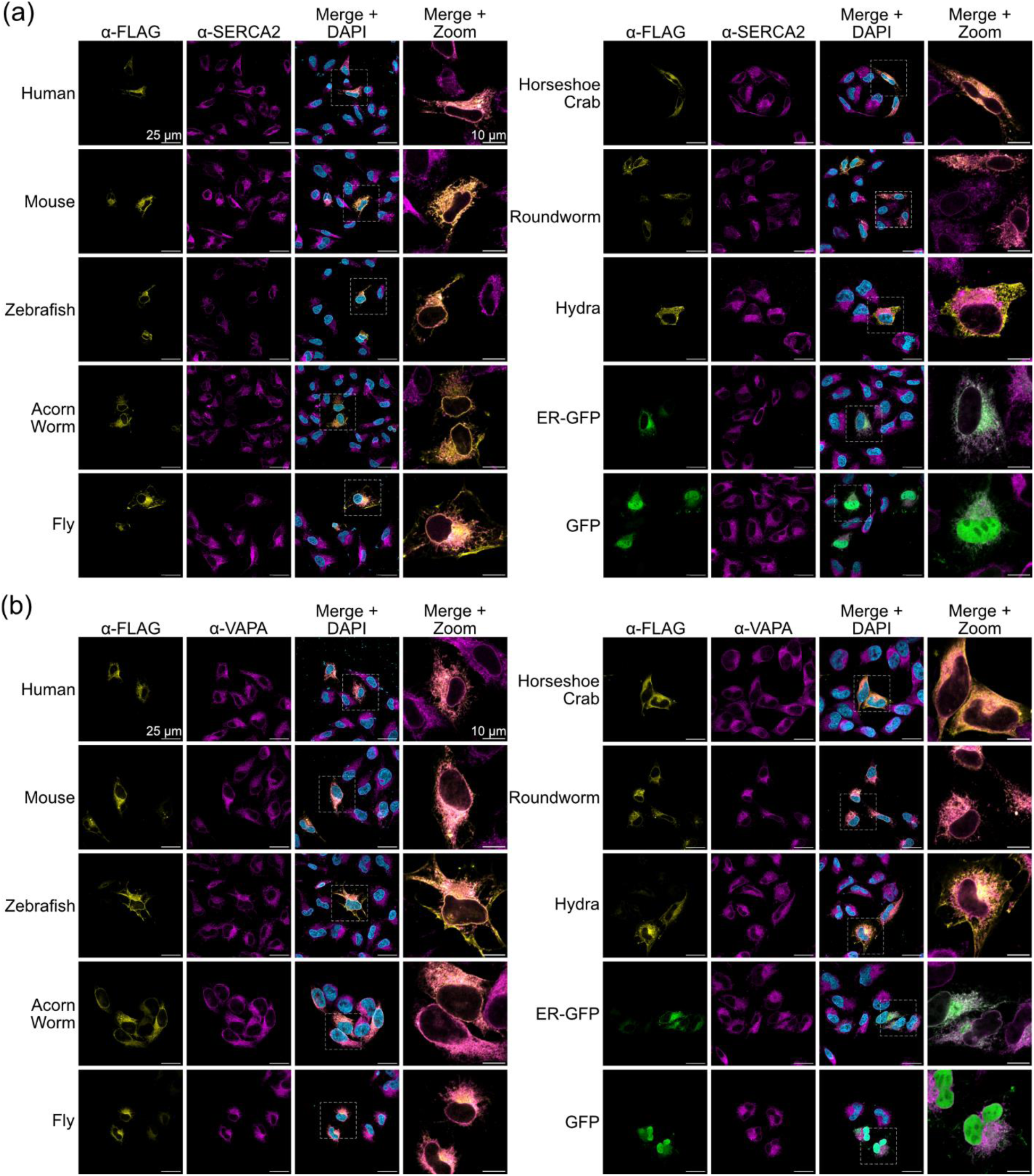
Representative confocal microscopy images of ANKLE2 orthologs.

**Figure S2:**
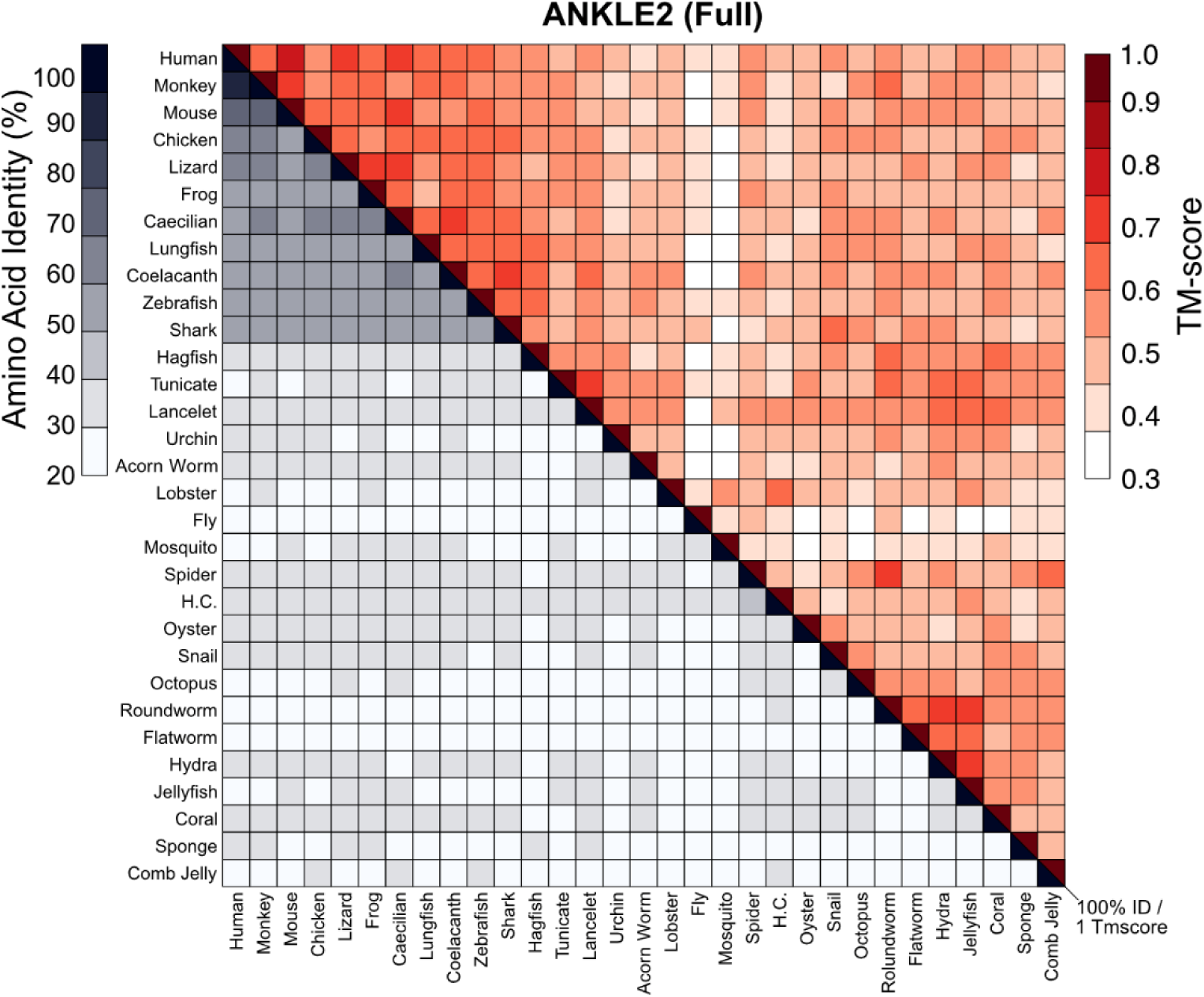
Sequence identity versus structural similarity matrix for full-length ANKLE2 orthologs.

**Figure S3:**
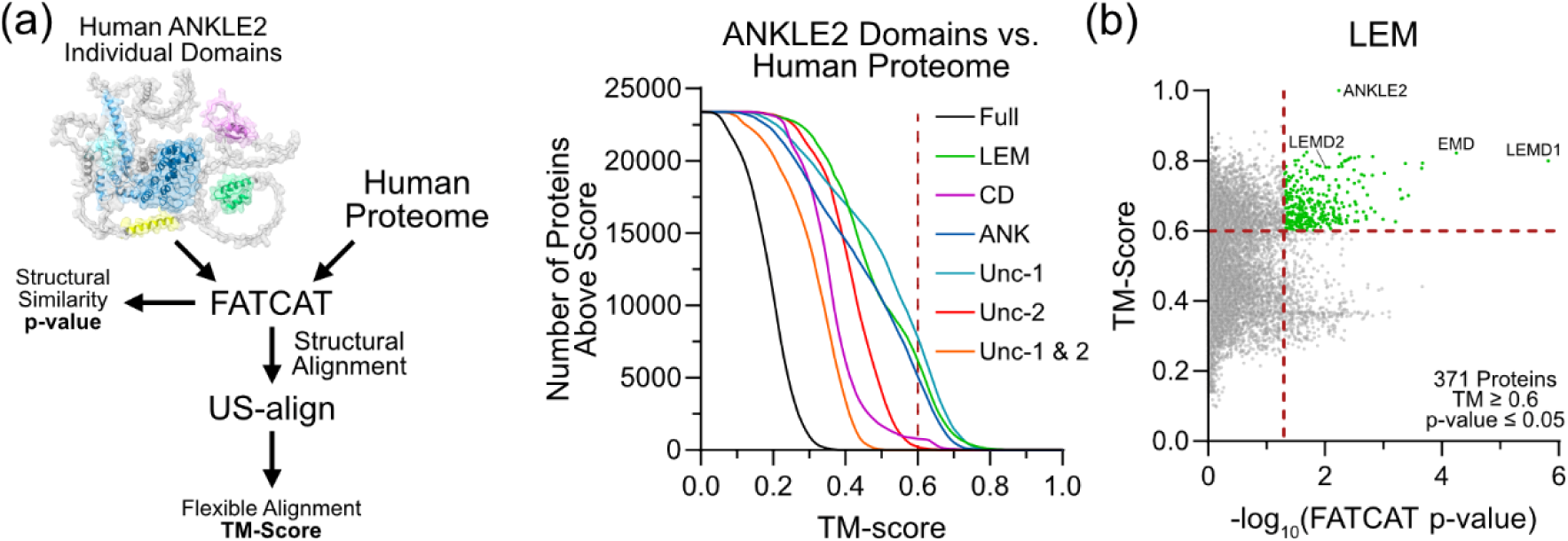
Foldseek and FATCAT/US-align as tools to evaluate similarity of individual ANKLE2 domains.

**Figure S4:**
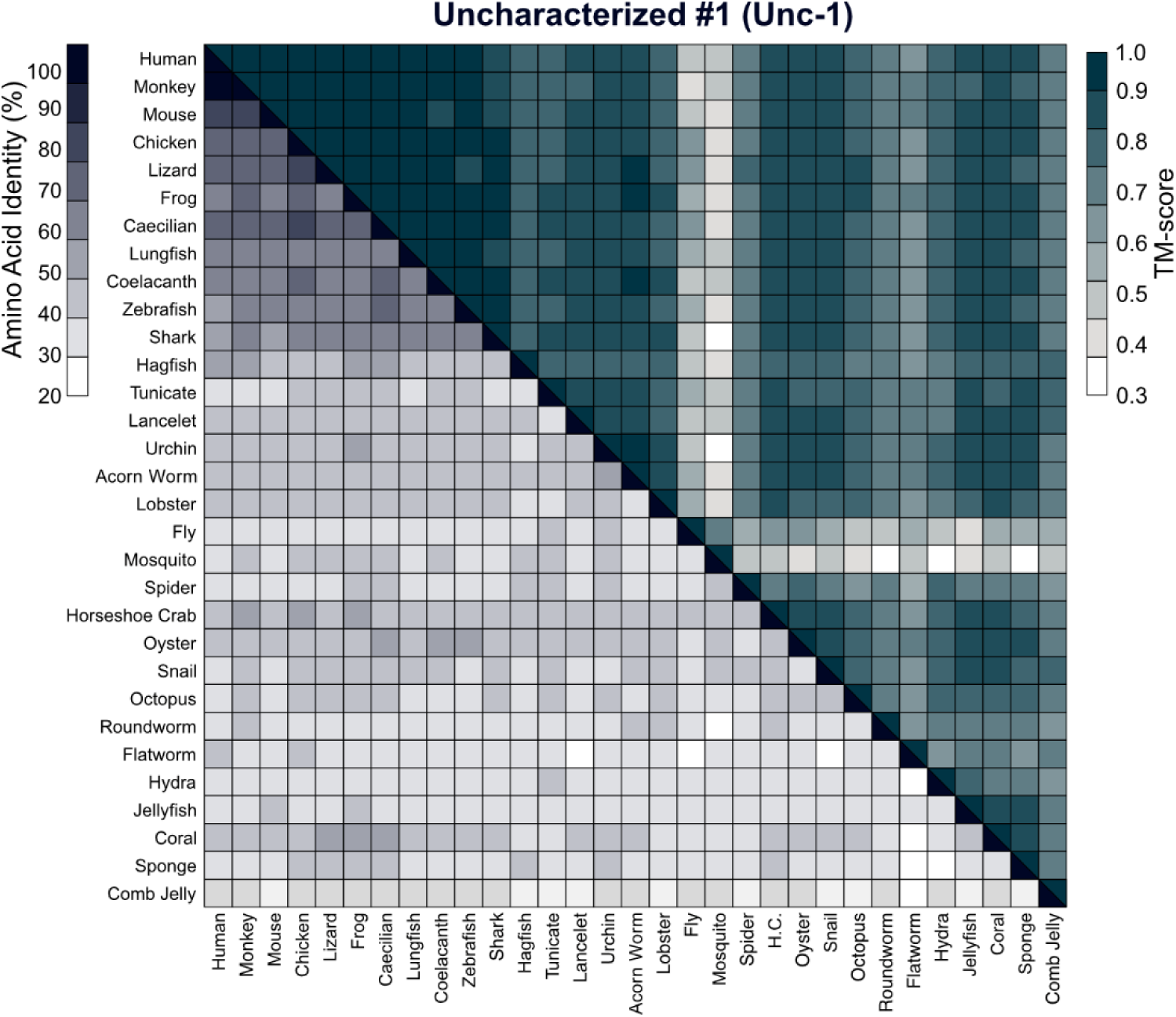
Sequence-structure matrix for uncharacterized domain #1.

**Figure S5:**
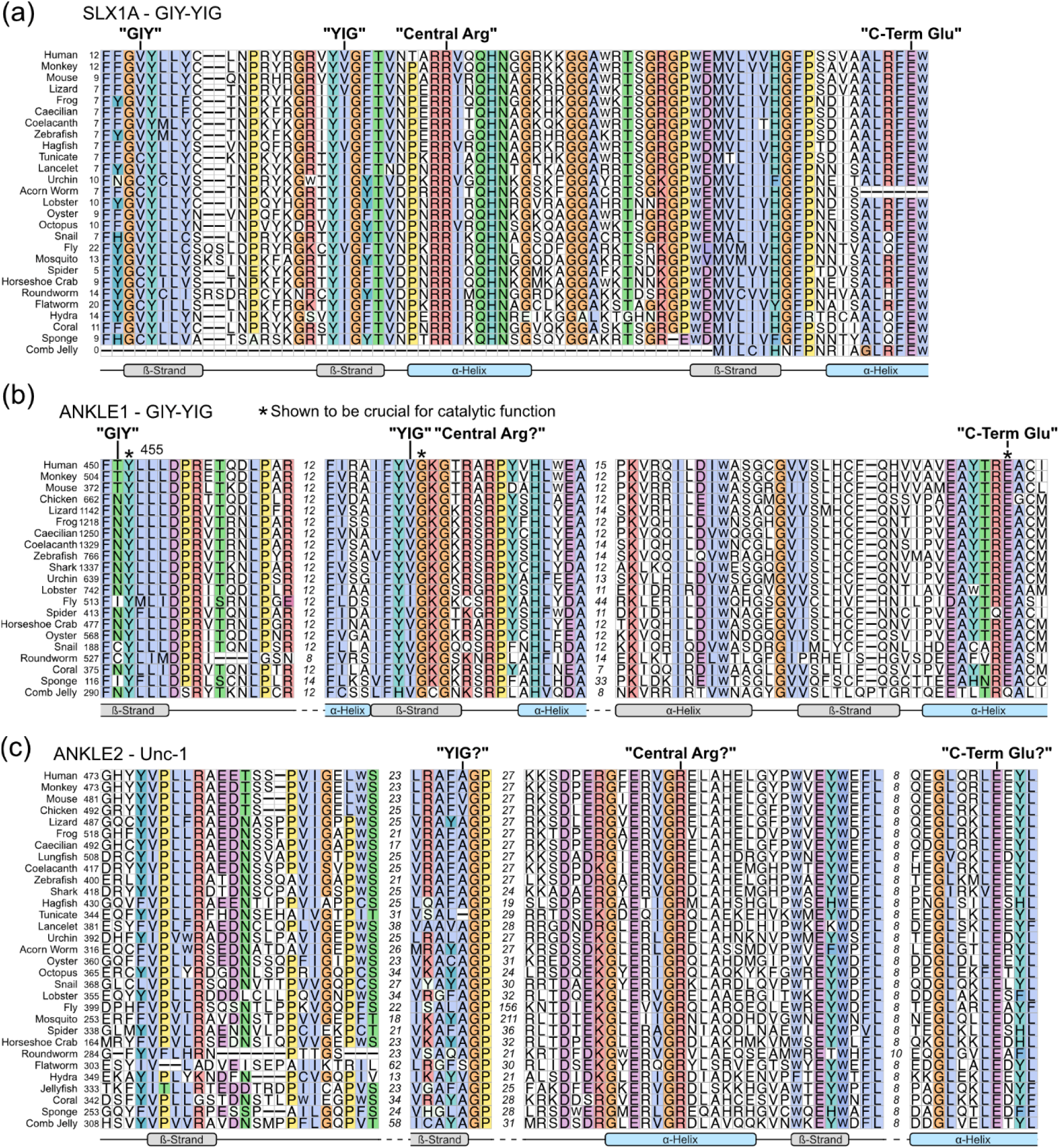
Sequence alignments of GIY-YIG regions of SLX1A, ANKLE1, and ANKLE2.

**Figure S6:**
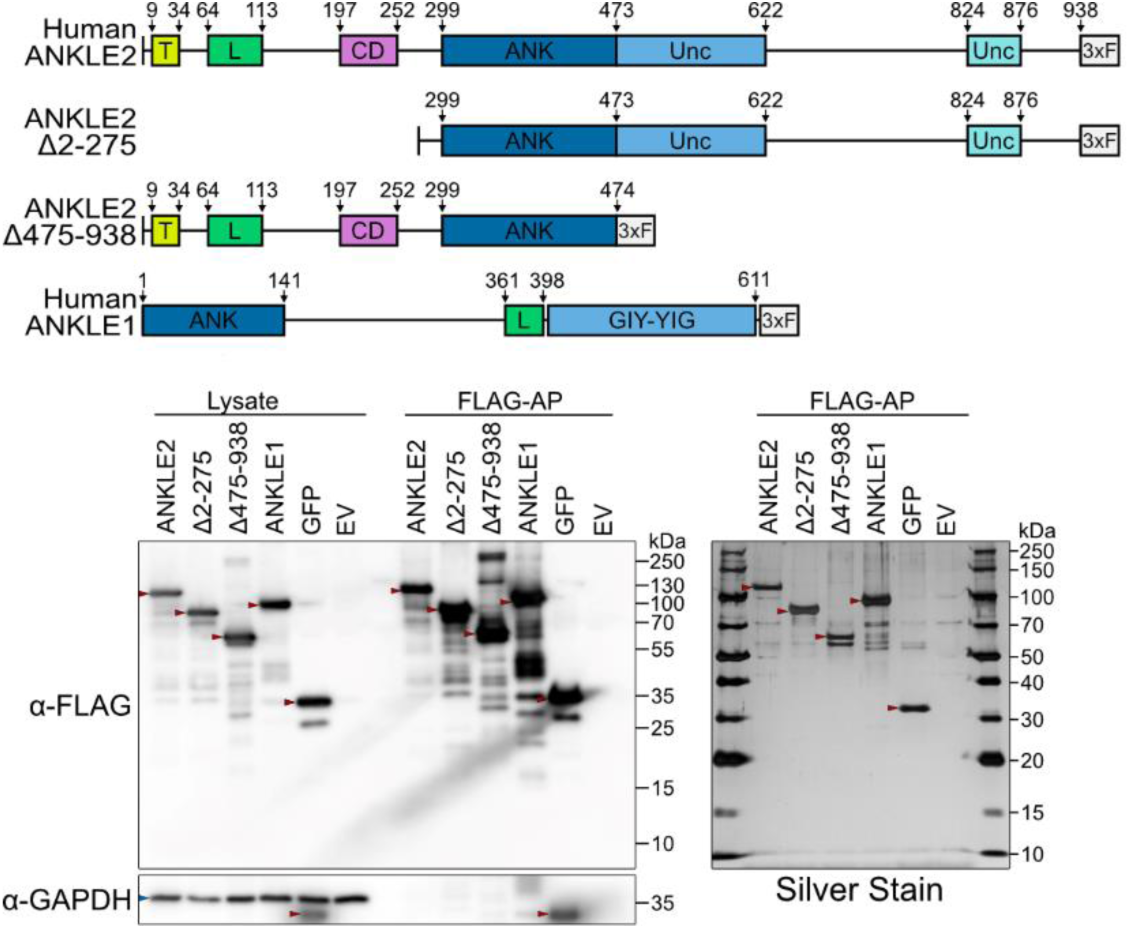
Protein production and purification validation.

**Figure S7:**
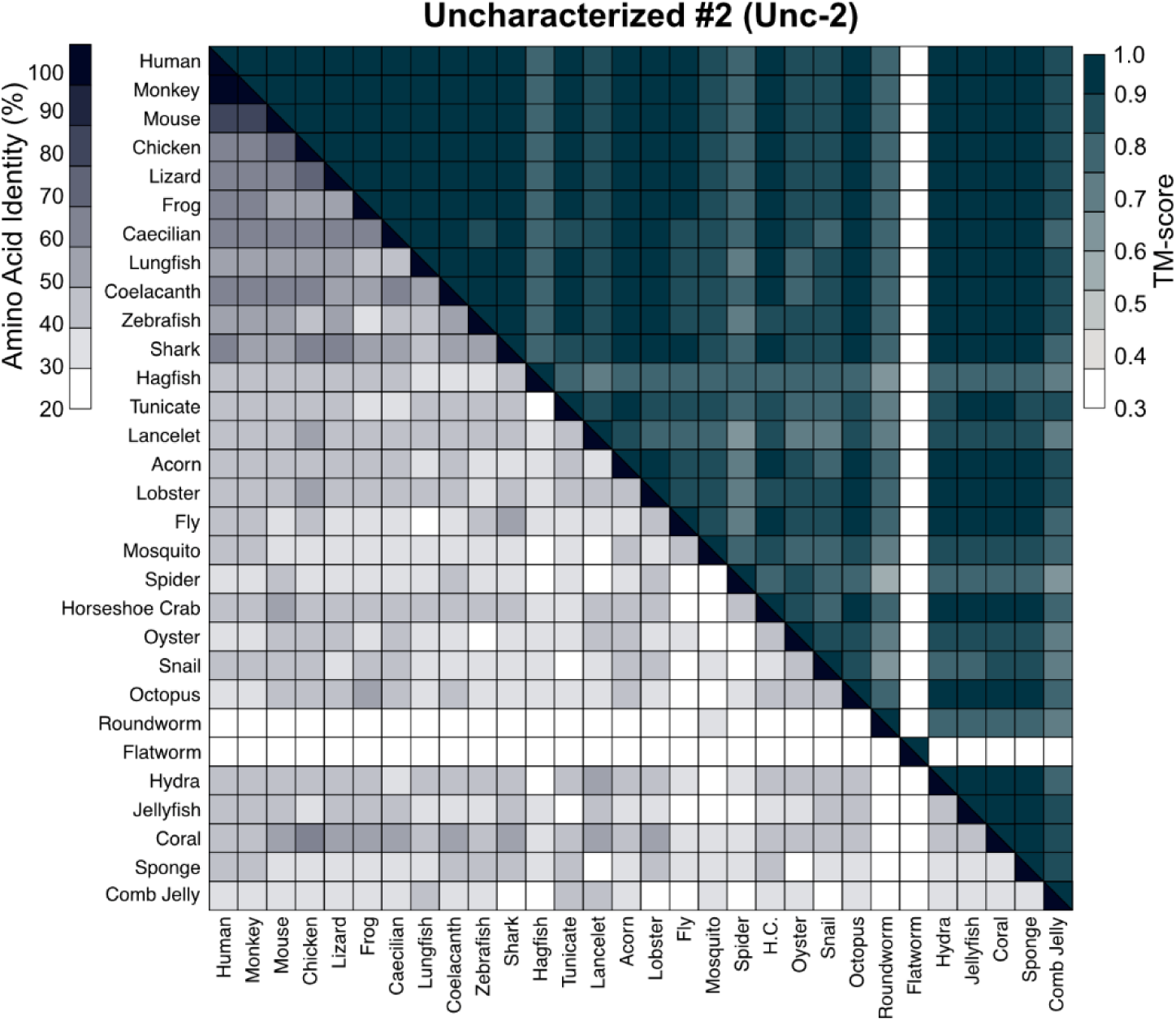
Sequence-structure matrix for uncharacterized domain #2.

**Figure S8:**
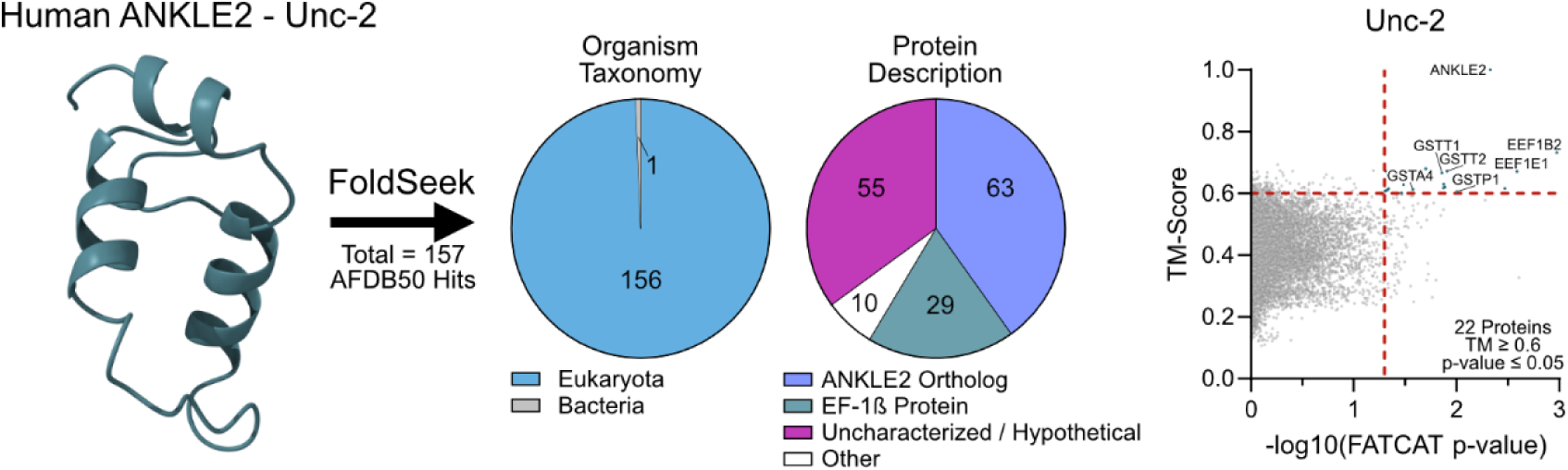
Foldseek results for ANKLE2 uncharacterized domain #2.

**Figure S9:**
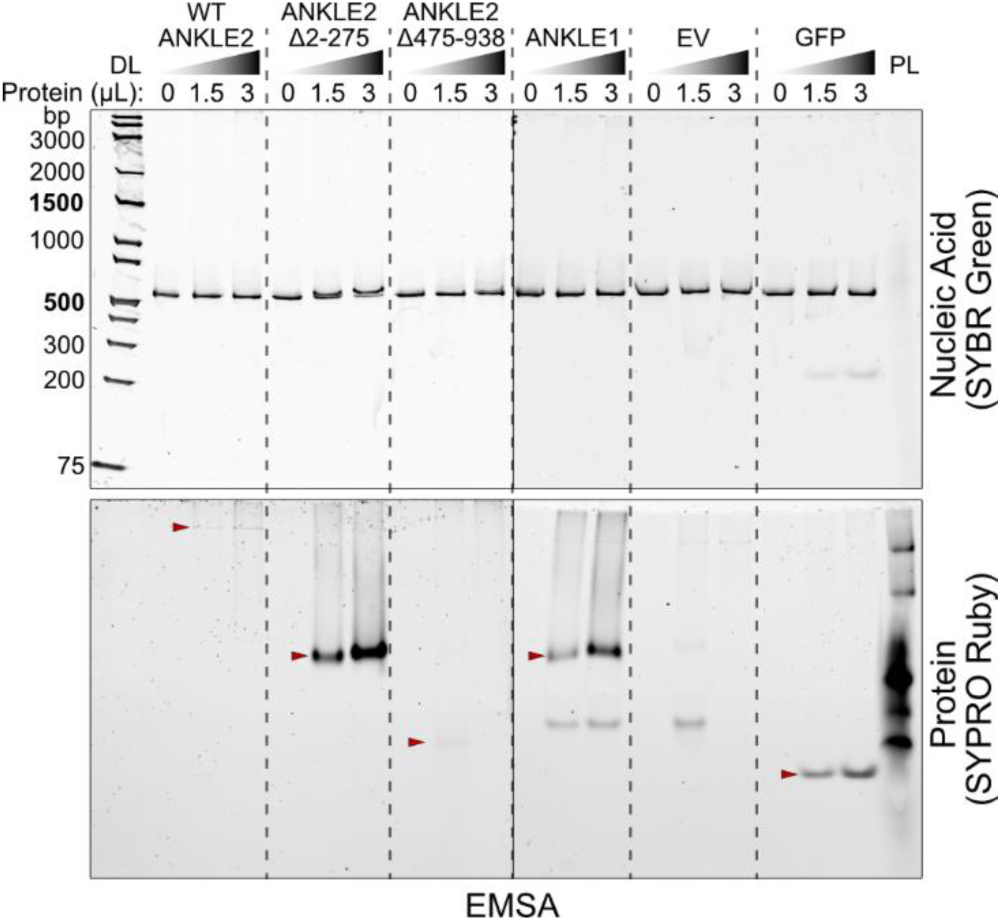
ANKLE2 does not bind with dsDNA by electrophoretic mobility shift assay.

**Figure S10:**
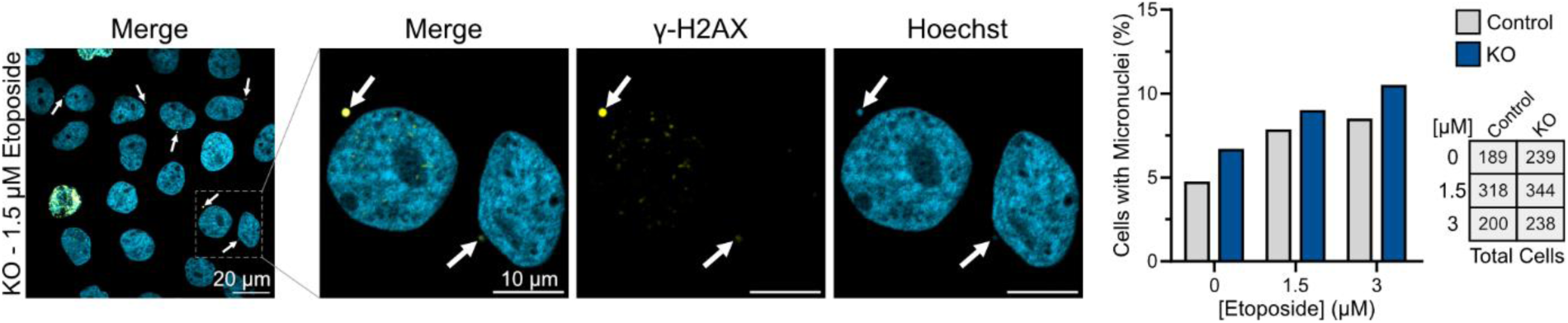
Observations of micronuclei during etoposide treatment.

**Table S1:** Foldseek, FATCAT, and US-align data for ANKLE2 ortholog comparisons and ANKLE2 domain vs. proteome wide searches.

**Table S2:**
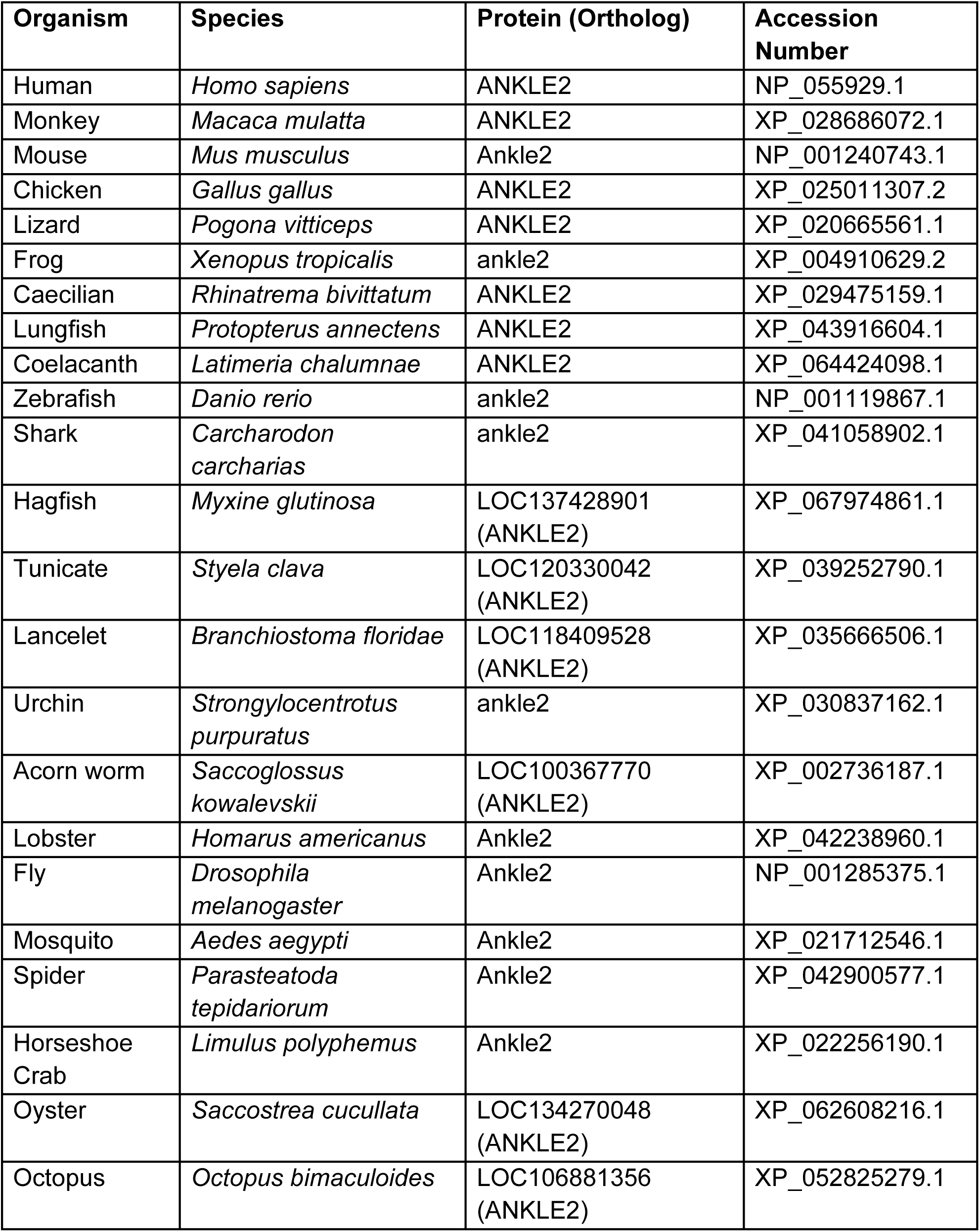
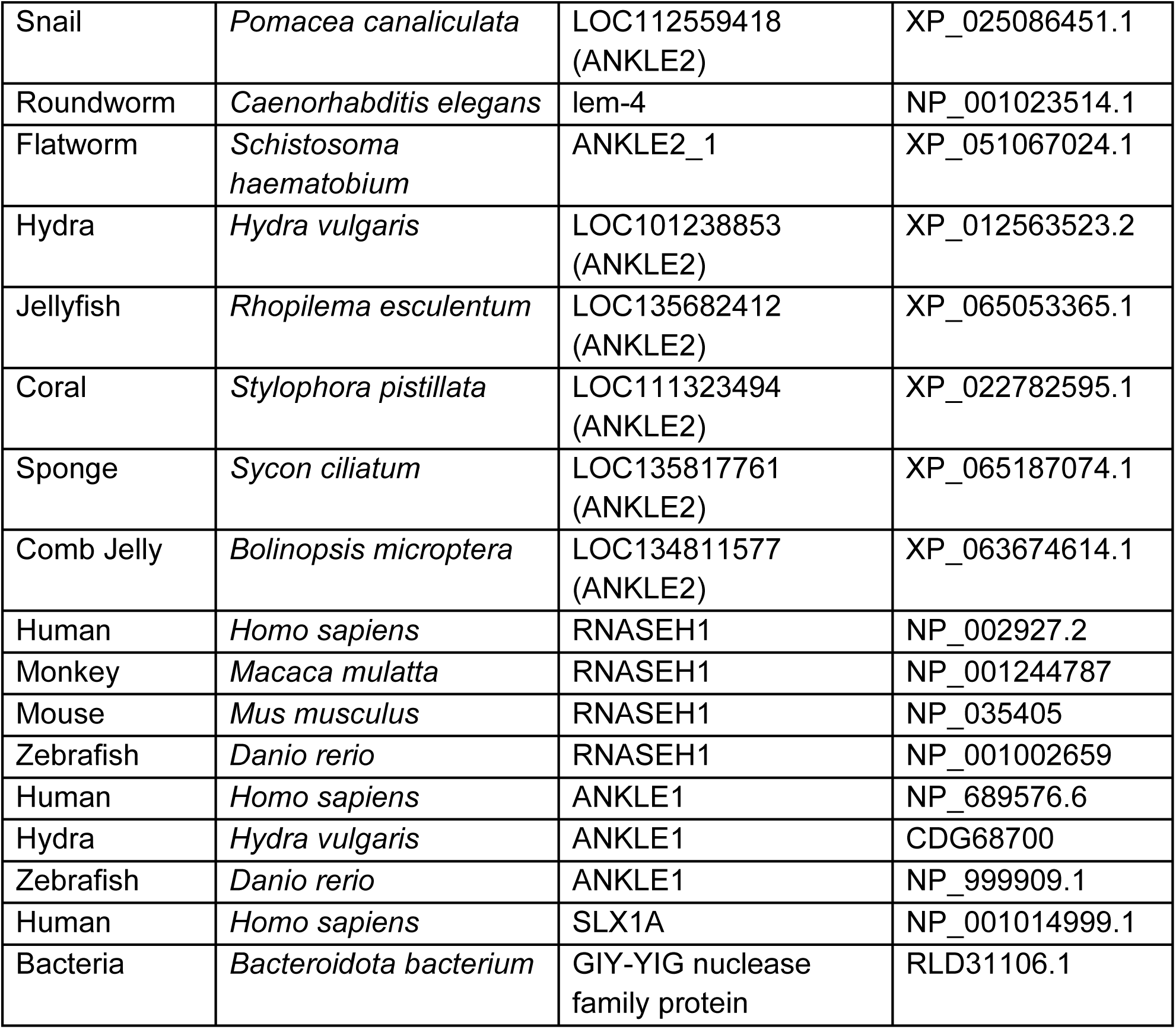
Sequence accession numbers for all protein sequences used.

**Table S3:**
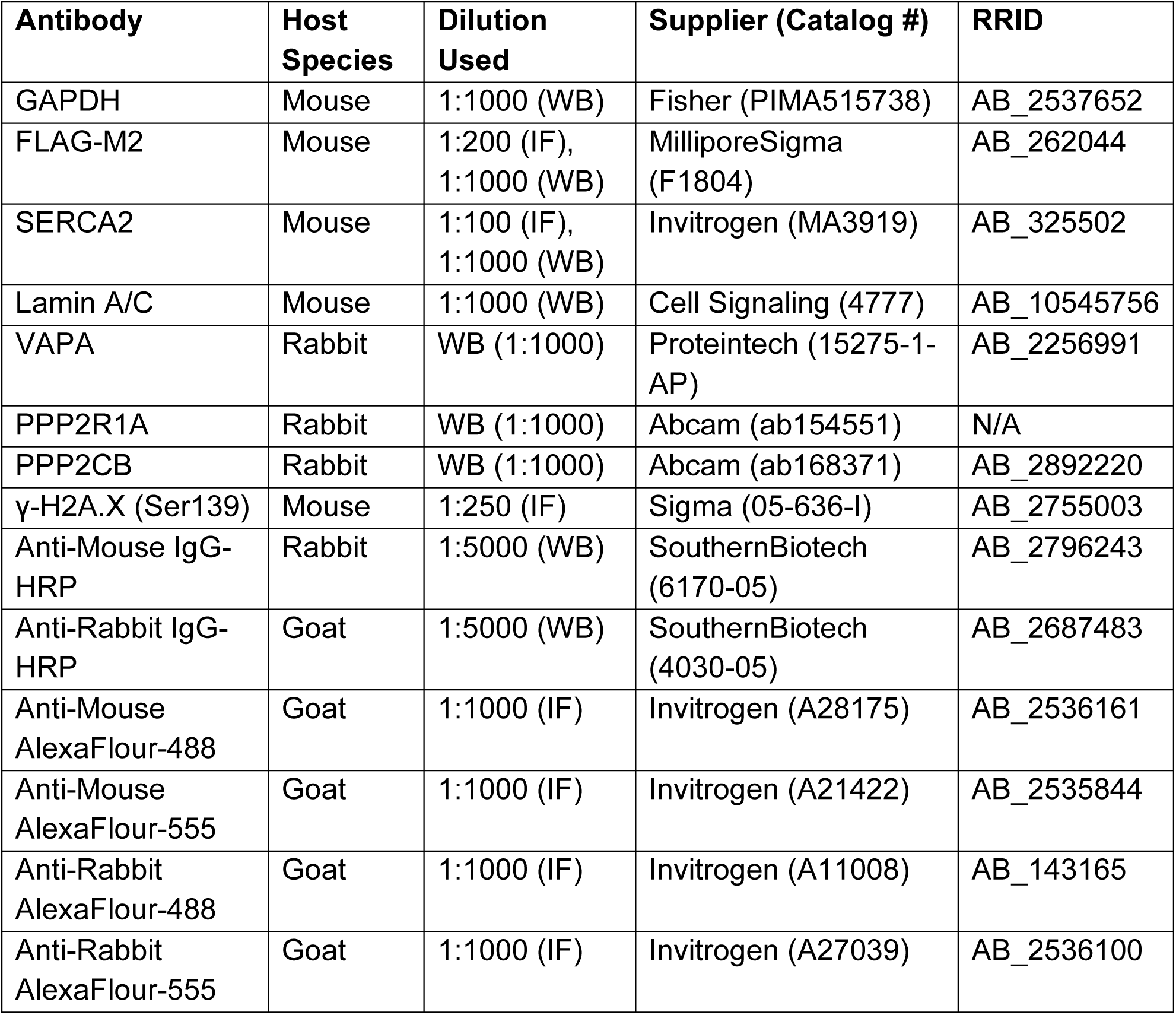
Antibody usage details for all antibodies used.

| **Antibody** | **Host Species** | **Dilution Used** | **Supplier (Catalog #)** | **RRID** |
| --- | --- | --- | --- | --- |
| GAPDH | Mouse | 1:1000 (WB) | Fisher (PIMA515738) | AB_2537652 |
| FLAG-M2 | Mouse | 1:200 (IF),<br>1:1000 (WB) | MilliporeSigma<br>(F1804) | AB_262044 |
| SERCA2 | Mouse | 1:100 (IF),<br>1:1000 (WB) | Invitrogen (MA3919) | AB_325502 |
| Lamin A/C | Mouse | 1:1000 (WB) | Cell Signaling (4777) | AB_10545756 |
| VAPA | Rabbit | WB (1:1000) | Proteintech (15275-1-AP) | AB_2256991 |
| PPP2R1A | Rabbit | WB (1:1000) | Abcam (ab154551) | N/A |
| PPP2CB | Rabbit | WB (1:1000) | Abcam (ab168371) | AB_2892220 |
| γ-H2A.X (Ser139) | Mouse | 1:250 (IF) | Sigma (05-636-I) | AB_2755003 |
| Anti-Mouse IgG-HRP | Rabbit | 1:5000 (WB) | SouthernBiotech<br>(6170-05) | AB_2796243 |
| Anti-Rabbit IgG-HRP | Goat | 1:5000 (WB) | SouthernBiotech<br>(4030-05) | AB_2687483 |
| Anti-Mouse AlexaFlour-488 | Goat | 1:1000 (IF) | Invitrogen (A28175) | AB_2536161 |
| Anti-Mouse AlexaFlour-555 | Goat | 1:1000 (IF) | Invitrogen (A21422) | AB_2535844 |
| Anti-Rabbit AlexaFlour-488 | Goat | 1:1000 (IF) | Invitrogen (A11008) | AB_143165 |
| Anti-Rabbit AlexaFlour-555 | Goat | 1:1000 (IF) | Invitrogen (A27039) | AB_2536100 |

